# A novel approach to localize cortical TMS effects

**DOI:** 10.1101/595603

**Authors:** Konstantin Weise, Ole Numssen, Axel Thielscher, Gesa Hartwigsen, Thomas R. Knösche

## Abstract

Despite the widespread use of transcranial magnetic stimulation (TMS), the precise cortical location underlying the physiological and behavioral stimulation effects are still only coarsely known. So far, mapping strategies rely on center of gravity approaches and therefore localize the stimulated cortical site only approximately and indirectly. Focusing on the motor cortex, we present a novel method to reliably determine the effectively stimulated cortical site at the individual subject level. The approach combines measurements of motor evoked potentials (MEPs) at different coil positions and orientations with numerical modeling of induced electric fields. We identify sharply bounded cortical areas around the gyral crowns and rims of the motor hand area as the origin of MEPs and show that the tangential component and the magnitude of the electric field is most relevant for the observed effect. To validate our approach, we determined motor thresholds for coil positions and orientations for the predicted cortical target. Our methods allows for the identification of optimal coil positions and orientations. Moreover, we used extensive uncertainty and sensitivity analyses to verify the robustness of the method and identify the most critical model parameters. Our generic approach improves the localization of the cortex area stimulated by TMS and may be transferred to other modalities such as language mapping.

## 1 INTRODUCTION

Transcranial magnetic stimulation (TMS) is capable of modulating motor and cognitive functions in the human brain. An important application of this technique is mapping structure–function relationships (see Bestmann and Feredoes, 2013; Sandrini et al., 2011; Siebner et al., 2009 for review). Answering the question of “*which part of the brain gets stimulated, and how do we know where it is?*” on the individual subject level is essential to understand brain physiology and structure function relationships. Such mappings are not only of fundamental neuroscientific interest, but also have practical clinical relevance, for example in the context of pre-surgical mapping for counseling and planning tumor resections or epilepsy surgeries. In a classical mapping study, the TMS coil is systematically moved over different positions and/or orientations while certain behavioral or physiological variables (e.g., the degree of speech impairment or the magnitude of motor evoked potentials) are measured (Picht, 2014). The coil position/orientation producing the strongest effect is then used as a proxy for the brain structures underlying the targeted effects, either in a simple way by direct projection onto the cortical surface (Krieg et al., 2014) or in a more sophisticated way by calculating the induced electric fields (Tarapore et al, 2013). However, this approach has some principal shortcomings. First, its capability to unambiguously determine the location and orientation of stimulated neural structures is limited: even if the coil configuration associated with the optimal effect can be found accurately, this coil configuration generates electric fields in a wide range of neural structures, such as radial cells in several parallel sulcal walls and tangential structures (e.g., axons) in gyral crowns. The field maximum does not reliably indicate which of these components is actually driving the effect. Second, as the search space has at least three dimensions (two for the position on the head surface, one for coil orientation), accurate mapping may require a very large number of stimulations to avoid undersampling. Consequently, previous studies remain controversial on fundamental aspects of the physiological TMS effects and their localization on the cortical surface. For instance, it is still unclear which part of the primary motor cortex is effectively stimulated by the TMS pulse (see Bungert et al., 2017; Laakso et al., 2018; Fox et al., 2004; Krieg et al., 2013), precluding strong conclusions on the cortical origin of the TMS effect. In particular, previous studies remain controversial with respect to the contribution of the gyral crowns and sulcal walls in the primary motor cortex (see Fox et al., 2004; Bungert et al., 2017; Opitz et al., 2013; Laakso et al., 2018; Krieg et al., 2013).

Resolving these limitations and establishing a link between coil position and location and size of affected cortical area in three dimensions (i.e., also in depth) is not trivial. It requires detailed knowledge of the electric field pattern at the individual level, biophysically motivated hypotheses on the mechanism of action by which the electric field causes neural excitation, and formal statistical testing to demonstrate the validity of the obtained results.

The induced electric field distribution strongly depends on several stimulation parameters such as intensity, location, and orientation of the TMS coil as well as the complex geometry of the individual brain (Thielscher et al., 2011), and several biophysical parameters, such as tissue conductivities and fractional anisotropy. Numerical modeling of the induced electric field is increasingly being increasingly used to address these issues (Bestmann, 2015; Thielscher et al., 2011; Thielscher et al., 2015), but has not become a standard procedure in medical and scientific applications so far. In particular, calculations based on subject-specific head meshes have improved our understanding of the impact of individual head anatomy on field distributions (Datta et al., 2010; De Lucia et al., 2007; Opitz et al., 2013; Opitz et al., 2011; Opitz et al., 2014; Thielscher et al., 2011). As such, numerical field calculations using anatomically detailed head models may assist the neurobiological interpretation of TMS effects, and aid the localization of the stimulated cortical area that underlies the observed physiological or behavioral effect (see Hartwigsen et al., 2015; Bungert et al., 2017). However, since a TMS pulse induces a distributed electric field over an extended part of the cortex, it is difficult to determine the location of neural activation, even if the field is computed in a reliable way.

Moreover, any approach that aims to make beneficial use of field models has to account for the uncertainties of the numerical simulations to yield accurate and robust conclusions. Uncertainties are particularly caused by the assumed ohmic tissue conductivities, which are only coarsely known. Such an approach also has to be further able to deal with uncertainties caused by the limited knowledge on how the induced field acts on different neuron types. Recently, field calculations have been combined with microscopic neural models based on accurate reconstructions from histology (Seo et al., 2017; Aberra et al. 2018). However, for now, validation of these models is still largely missing to a large extent and the conclusions strongly depend on model details such as the types of the included neural elements. Consequently, the results of previous studies strongly differ and do not provide reliable conclusions yet.

In this study, we introduce a novel TMS mapping approach that links biophysical modeling of the induced electric field with physiological measurements within a principled statistical testing framework to determine the stimulated cortical area on the individual subject level. Our approach is based on the assumption of a unique functional relationship between the observed physiological TMS effect and the electric field induced at the cortical location underlying this effect. Given an experimental effect that linearly or non-linearly scales with stimulation intensity, one can assume that this effect also scales with the degree of excitation of the specific neuronal population, which is functionally linked to it. Hence, the functional relationship between the electric field component that coincides spatially and orientation-wise with this population and the observed effect should be invariant across experimental conditions, that is, different orientations and positions of the TMS coil. Consequently, the stimulated cortical area can be localized by determining the brain area in which the induced field shows a clear functional relationship between the measured effects across conditions. Note that this area does not have to coincide with the field maximum. Similar, but more restricted, localization approaches were used in previous studies. For instance, targeting the hand area of the primary motor cortex, Bungert et al. (2017) employed a statistical approach based on the experimentally determined motor thresholds at different coil orientations. Laakso et al. (2018) used a similar strategy, but investigated the influence of different coil positions while keeping the orientation constant. These studies demonstrate the principal validity of the rationale to localize the stimulated cortical area using the functional relationship between calculated fields and the observed effects. However, they remain restricted in several important aspects. First, the ability for a precise functional localization at the single-subject level was not demonstrated. Second, it remains unknown how many experimental conditions are needed to achieve a satisfying localization result and how these coil positions and orientations should be chosen. Third, the robustness of the approaches to uncertainties of tissue conductivities was not examined. Finally, all of the aforementioned publications lack experimental validation.

Our novel approach differs from these prior studies in two important aspects to advance localization at the single-subject level: 1) it combines multiple stimulations with different coil positions and orientations. 2) Instead of relying on motor threshold, it exploits entire input–output curves (I/O curves; relationship between stimulation intensity and MEP amplitude; see Fig. 1). We show that our method provides a means of precisely localizing the individual cortical area that is responsible for the observed motor output. We prove its stability and robustness using comprehensive permutation tests and a rigorous uncertainty and sensitivity analysis based on the generalized polynomial chaos (gPC) approach (Le Maitre et al., 2010). To account for the limited knowledge on the neural target structures of TMS, we tested several components of the induced field, and consistently found the tangential field component and the field magnitude to be the relevant quantities for modulating the observed effect. Importantly, we demonstrate that unique results can be obtained with relatively few measurements and indicate how the respective coil positions and orientations should be chosen. We validated our method by numerically optimizing the individual TMS coil position and orientation to effectively stimulate the identified cortical targets. These coil positions and orientations were shown to produce lower MTs than any other tested coil configuration. Our approach improves the localization of effectively stimulated areas during TMS. While demonstrated for the motor cortex, our approach is generic and can be applied to mapping procedures in other domains such as language.

**Figure 1:**
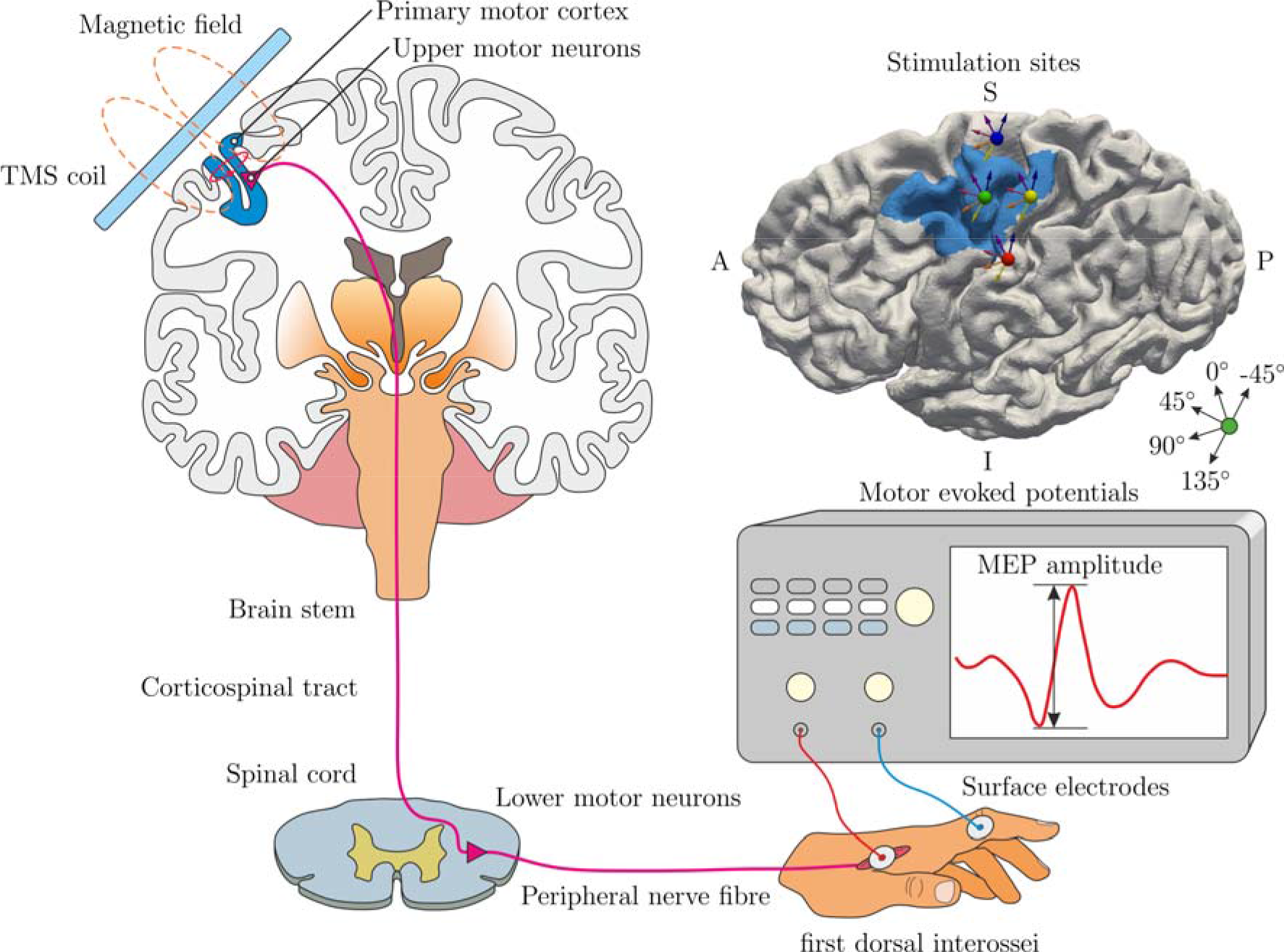
Schematic representation of the experimental procedure of the TMS experiments. Top left: The TMS coil is located tangentially to the skull over the primary motor cortex (M1). A time changing current in the coil generates a time changing magnetic field, which induces an electric field in the brain. This depolarizes the upper motor neurons with corticospinal efferents. Bottom: Action potentials from the upper motor neurons excite the lower motor neurons in the spinal cord, evoked action potentials travel through the peripheral nerves to the first dorsal interossei (FDI) of the hand. Sum potentials (motor–evoked potentials, MEP) are recorded from hand muscles using a classical belly tendon montage, i.e. between the dorsal interosseus and proximal interphalangeal joint of the index finger. Top right: example cortical surface with region of interest (blue), showing positions of the coil centers (colored spheres) and the coil orientations (arrows) for the 20 experimental conditions.

## 2 MATERIALS AND METHODS

We developed a novel framework to localize the neuronal populations that are responsible for effects of TMS by combining behavioral responses with numerical modeling. We applied this to primary motor cortex stimulation and electrophysiological measurements of muscle activation. First, we describe the experimental design to elicit and measure MEPs (2.1). Second, we show how to calculate the TMS induced electric field distribution in the subjects’ heads (2.2). This covers models for the head, the TMS coils, as well as the differential equations numerically solved to determine the electric field inside the brain. Third, we present a new measure to quantify the correlation between the induced electric field and the behavioral or physiological stimulation effect, the *congruence factor*. Locations with high congruence are likely to house neural populations that are causally linked to the observed MEP (2.3). We then show how the results are analyzed in terms of their sensitivity towards uncertain model parameters such as the electrical conductivities of the brain tissues and the measured MEPs using a generalized polynomial chaos (gPC) approach (2.4). Finally, the validation procedure for our results is outlined (2.5).

### 2.1 TMS Experiments

The experimental setup is shown in Fig. 1. Fifteen healthy, right-handed participants (seven female, age 22-34 years) with a mean laterality index of 92.93 (SD = 10.66) according to the Edinburgh Handedness Inventory were recruited. Subject inclusion was in accordance with the published safety guidelines on patient selection for TMS studies (Rossi et al., 2009; Rossini et al., 2015). Written informed consent was obtained from all participants prior to the examination. The study was performed according to the guidelines of the Declaration of Helsinki and approved by the local Ethics committee of the Medical Faculty of the University of Leipzig.

Stimulation was applied with a MagPro X100 stimulator (MagVenture, firmware Version 7.1.1) and CB-60 figure-of-eight coils, guided by a neuronavigation system (software: Localite, Germany, Sankt Augustin; camera: Polaris Spectra, NDI, Canada, Waterloo).

MEPs were recorded from the subjects’ right hand index finger with one surface electrode positioned over the muscle belly of the FDI and one at the proximal interphalangeal joint (PIP). The electrodes were connected to a patient amplifier system (D-360, DigitimerLtd., UK, Welwyn Garden City; bandpass filtered from 10 Hz to 2 kHz), which in turn was connected to a data acquisition interface (Power1401 MK-II, CED Ltd., UK, Cambridge, 2 kHz sampling rate). Stimulation control and recording was performed with Signal (CED Ltd., version 4.11).

Localization of the MEP stimulation hotspot was guided by individually transformed M1 coordinates based on the standardized group coordinates from a meta-analysis (Mayka et al. 2006). These coordinates were transformed to the individual subject’s space by using the inverse of the normalization transformation in SPM (Penny et al., 2007; https://www.fil.ion.ucl.ac.uk/spm/). The individual MEP producing hotspot M1_45°_ and its corresponding resting motor threshold (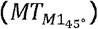) were determined. 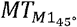 was defined as the minimum stimulation intensity, which evokes MEPs with an amplitude of at least 50 μV in at least 5 out of 10 consecutive stimulations (Pascual-Leone and Torres, 1993; Rothwell et al., 1999; Conforto et al. 2004).

In relation to M1_45°_, five additional conditions, shown in Fig. 2(a), with different stimulation sites and coil orientations (see Fig.2a) were defined in the following way: in Experiment I, the TMS coil was located over M1 and 2 cm posterior (P). At both sites, three coil orientations with respect to M1_45°_ were investigated, namely M1_0°_/P_0°_ (−45° from M1_45°_), M1_45°_/P_45°_, and M1_90°_/P_90°_ (+45° from M1_45°_), resulting in six experimental conditions.

**Figure 2:**
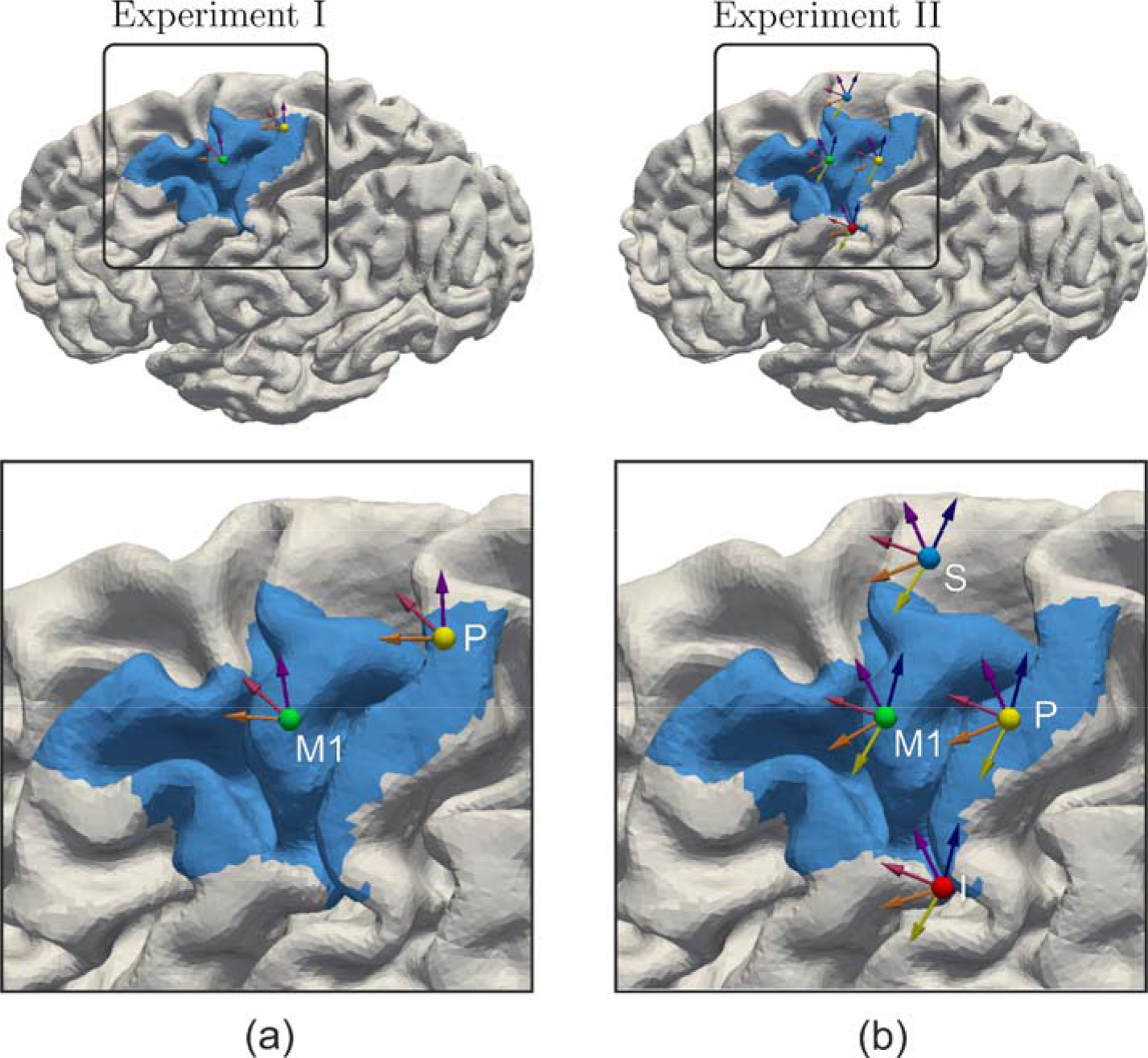
Coil positions and orientations used in (a) Experiment I and (b) Experiment II. The locations are exemplarily shown for subject S8. The number of experimental conditions increases from six to 20. The blue area is the region of interest, comprised of the somatosensory cortex S (BA 1, BA 3), M1 (BA 4), and the dorsal part of the premotor cortex (BA 6).

Experiment II included three of the subjects from Experiment I and the number of conditions was increased to further investigate the influence of different coil positions and orientations on the determination of the effective stimulation site. In addition to M1 and P, two more coil positions were included, shown in Fig. 2(b), 2 cm inferior and 2 cm superior to M1, respectively (Fig.2b). For each position, the number of coil orientations was increased to 5 (− 90°, −45°, 0°, 45°, 90°) with respect to M1_45°_, resulting in 20 experimental conditions.

In both experiments, single biphasic pulses with an inter stimulus interval of 5 s (Experiment I) or 4 s (Experiment II) were applied for each condition. The coil positions/orientations were recorded by the neuronavigation system. The MEPs were lowpass filtered with a 6^th^ order Butterworth filter with a cutoff frequency of 500 Hz. Afterwards, the peak-to-peak amplitudes of the MEPs were calculated in a time window of 18 to 35 ms after the TMS pulse (see Fig 3a for an example MEP). Stimulation intensities were chosen to sample the complete I/O curve for each experimental condition, unless maximal stimulator output was reached before (Fig. 3(b)). Intensity was increased in steps of 2% stimulator output (MSO), or 1% respectively for intensity ranges of high I/O gradients (cf. Bungert et al., 2017). For each intensity, 3-5 stimulations were performed to determine an average MEP amplitude. Trials with deviations in ± ±coil position of ±3 mm and orientation of ±5° with respect to each axes were removed. A typical set of data points is shown in Fig. 3(b) (blue dots). Thereafter, a sigmoidal function was fitted in a least-square sense:

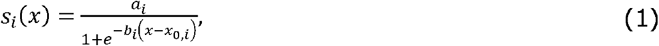

where *a*_*i*_ is the saturation amplitude, *b*_*i*_ the slope, and *x_0,i_*, is the location of the turning point on the abscissa. If only a part of the I/O curve could be determined experimentally, a sigmoidal function could not be reliably fitted and an exponential or linear function was used instead. The selection of the optimal model was performed using the Akaike information criterion (AIC, Akaike, 1974). The procedure was repeated for all experimental conditions (i.e., for different coil positions and orientations) in a pseudo-randomized order.

**Figure 3:**
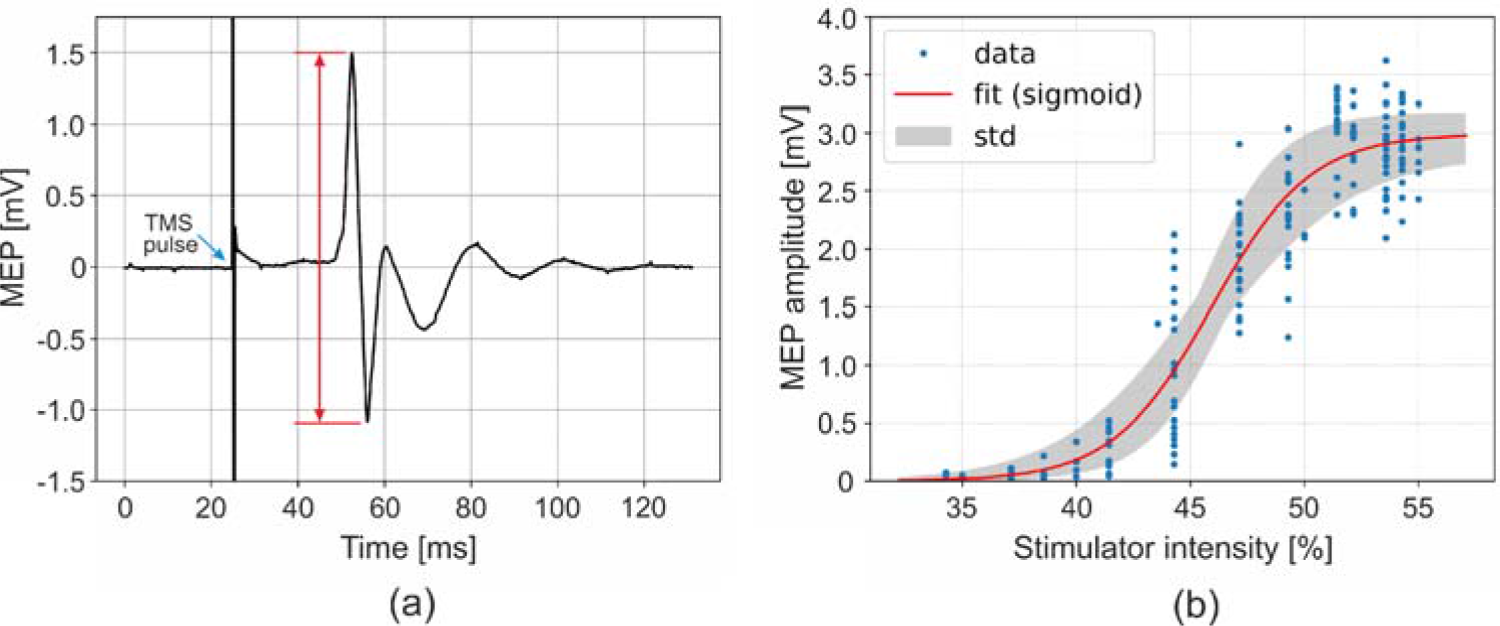
(a) Measured motor evoked potential (MEP) showing the stimulation artifact (blue arrow) and highlighting the peak-to-peak amplitude (red arrow). (b) I/O curve characterizing the MEP amplitudes as function of the stimulator intensity for one experimental condition. Blue dots: MEP amplitudes for the different stimulator intensities. Red curve: fitted analytical function. Depending on the best Akaike information criterion (AIC), MEP amplitudes were fitted to sigmoidal, exponential or linear functions (presented example: sigmoid function). Black lines: possible I/O curves resulting from uncertainty in the experimental data.

### 2.2 Numerical simulations of the induced electric field

The calculations of the electric field were conducted with SimNIBS v2.0 (Thielscher et al., 2015) using high-resolution anisotropic finite element models (FEMs), as exemplary shown for one subjects in Fig. 4. The individual head models were generated from MRI data using the pipeline described in Windhoff et al. (2013), employing FreeSurfer (http://surfer.nmr.mgh.harvard.edu/, Dale et al., 1999; Fischl et al., 1999) and FSL (https://fsl.fmrib.ox.ac.uk/fsl/fslwiki/FSL, Woolrich et al., 2009; Smith et al., 2004; Jenkinson et al., 2012). The head models were composed of ~1.3·10^6^ nodes and ~7·10^6^ tetrahedra. T1 and T2 images were used for segmenting the main tissues of the head: scalp, skull, grey matter (GM), white matter (WM), and cerebro-spinal fluid (CSF). Diffusion weighted images were used to reconstruct the conductivity tensors in the WM using the volume normalized mapping approach (Güllmar et al., 2010). To this end, the following structural images were acquired with a 3 Tesla MRI scanner (Siemens Verio or Skyra) and a 32 channel head coil. The following images were acquired: (i) T1-weighted: MPRAGE with 176 sagittal slices, matrix size = 256 × 240, voxel size = 1 × 1 × 1mm^3^, flip angle 9°, TR/TE/TI = 2300/2.98/900ms (Repetition, Spin echo, Inversion Time), (ii) T2-weighted: 192 sagittal slices, matrix size = 256 = 258, voxel size = 0.488 × .488 × 1mm³, flip angle 120° TR/TE = 5000/395ms (iii) diffusion MRI (67 axial slices, matrix size 128 × 128, voxel size 1.71875 × 1.71875 × 1.7mm^3^, TE/TR 80/7000ms, flip angle 90°, 67 diffusion directions, b-value 1000s/mm³^3^. An additional b0 image with reversed encoding direction was recorded for distortion correction with FSL topup and eddy. The T1 image was also used for neuronavigation during TMS. If adequate scans (age less than 1 year) already existed for the subjects in the image database, these scans were utilized.

**Figure 4:**
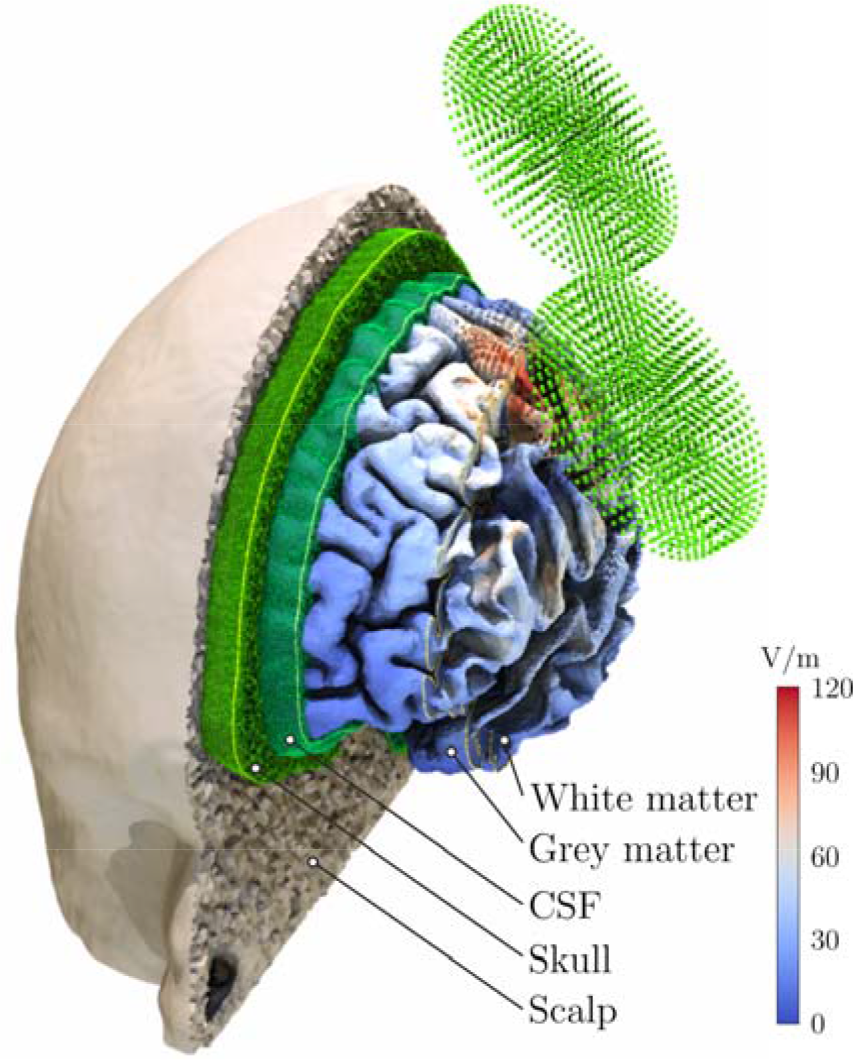
Example of the realistic anisotropic head model of one subject, used for the numerical simulations of the induced electric field. The model consists of 1.26 · 10^6^ nodes and 7.12 · 10^6^ tetrahedra. The TMS coil is modeled using 4440 magnetic dipoles (green spheres) with optimized dipole moments located in five layers. The grey matter surface is color coded with the magnitude of the induced electric field for 1 A/μs intensity.

All TMS coils were individually modelled by magnetic dipoles based on X-ray images. Coil wiring differences (shifts of several millimeters and tiltings of about 2-5°) were observed and accounted for. Each coil model consisted of ~4500 magnetic dipoles. These dipole models were compared to a detailed current density based FEM model using Comsol Multiphysics v4.4 (COMSOL, Inc., Burlington, MA, USA) and yielded magnetic field errors of < 0.1% at a distance of 15 mm.

The magnetic field produced by the coil was calculated in advance in terms of the magnetic vector potential *A*. The primary electric field is then given by *E*_*p*_ = −jωA, where ω = 2π*f* is theangular frequency of the biphasic TMS pulse. The electric potential *ϕ* in the nodes is calculated by solving the Laplace equation *∇* . ([σ]*∇*ϕ) = 0, considering anisotropic *∇* · ([σ]**∇**ϕ) = 0 conductivity tensors [*σ*] inside each element together with the boundary conditions given by the law of current conservation **∇* ·j = 0*. After calculating the secondary electric field *E*_*S*_ = −*∇*ϕ, the total induced electric field is given by *E* = −*jωA* − *∇*ϕ. The conductivity values for the five examined tissues (σ_*Scalp*_ = 0.465 S/m, σ_*Skull*_ = 0.01 S/m, σ_*GM*_ = 0.275 S/m, σ_*WM*_ = 0.126 S/m, σ_*CSF*_ = 1.654 S/m) were taken from Thielscher et al. (2011) and Wagner et al. (2004). A more detailed description about the FEM solver is given in Windhoff et al. (2013, supplemental material).

Individual coil positions and orientations relative to the subject’s head were saved by the neuronavigation system for each stimulation. These coil configurations were used for electrical field calculations in SimNIBS. A region of interest (ROI) was defined covering the somatosensory cortex (BA 1, BA 3), M1 (BA 4), and the dorsal part of the premotor cortex (BA 6) with Freesurfer and a mask was created for the Freesurfer average template and transformed to each individual subject’s brain.

The following analyses were performed on the midlayer between the outer surfaces of the GM and WM compartments in order to avoid boundary effects of the electric field due to conductivity discontinuities. The electric field was interpolated in the nodes of this surface in a post-processing step. Fig. 5 shows the magnitude |*E*|, the normal component |*E*_┴_| and the tangential component |E_‖_| of the electric field at the midlayer surface for three different coil positions and orientations in one exemplary subject. The different electric field distributions are shown in the highlighted ROI for the different experimental conditions.

**Figure 5:**
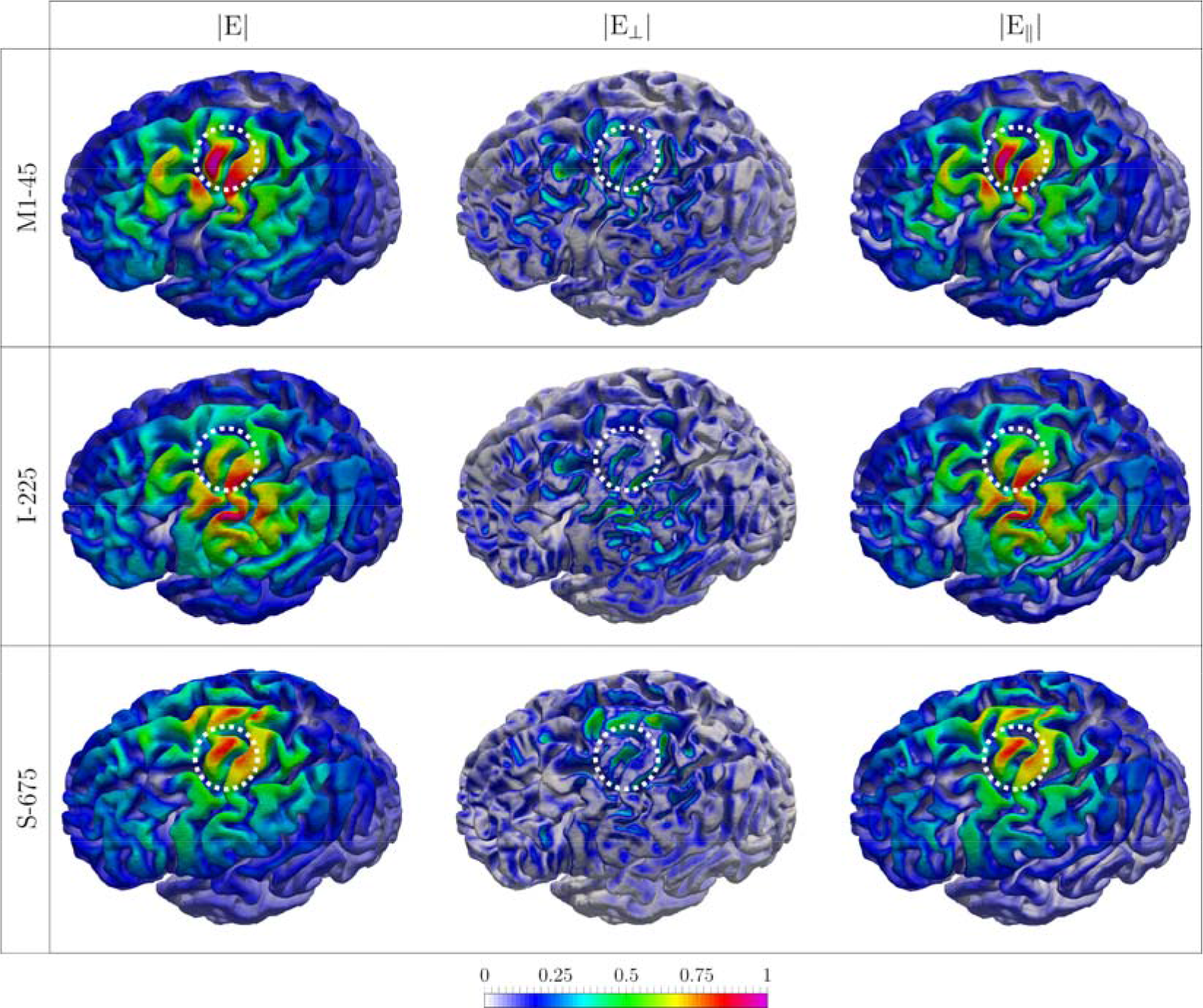
Three electric field distributions, i.e. the magnitude of the electric field |*E*|, the normal component |*E*_┴_|, and the tangential component |E_‖_| for one exemplary subject. The electric field was normalized to allow comparability between the different experimental conditions. The white circle shows the M1 hand knob area.

### 2.3 Determining the site of stimulation

The core concept of the proposed method is illustrated in Fig. 6. We assume that at the site of activation the correlate between electric field and MEP is stable, i.e. the same electric field evokes the same behavioral output independent of the location or orientation of the TMS coil. Exploiting this property, we can single out the site of stimulation can be singled out by calculating *cortical I/O curves* that represent the relationship between the electric field in the cortex and the resulting MEP, and then comparing the cortical I/O curves of the different experimental conditions. At the true cortical site of stimulation, the I/O curves of all conditions should be similar. Practically, this was achieved by transforming the measured I/O curve for each condition (representing the relationship between stimulation intensity in percent of maximal stimulator output, %MSO, and MEP amplitude) to E-MEP-curves (representing the relationship between the electric field at a particular cortex location and the respective MEP amplitude). The electric field distribution throughout the brain was computed as a function of the stimulation intensity using the numerical techniques described above, thereby taking 100% µMSO as corresponding to a maximal change of the coil current of 140 A/μs for the used stimulator-coil combination. Due to the linear relationship between electric field strength and stimulator intensity, the E-MEP curves were shifted and horizontally scaled versions of the measured I/O curve, with different shift and scale parameters in each position. Hence, the function types of each I/O curve and their corresponding E-MEP curves are similar (i.e. sigmoidal, exponential, or linear). E-MEP curves can be determined for all different components of the electric field vector (|*E*|, |*E*_┴_|, |E_‖_|) or, in principle, any other derived quantities thereof. This approach allows computing a position-wise *congruence factor c*(*r*), which quantifies the similarity between the E-MEP curves of the different experimental conditions.

**Figure 6:**
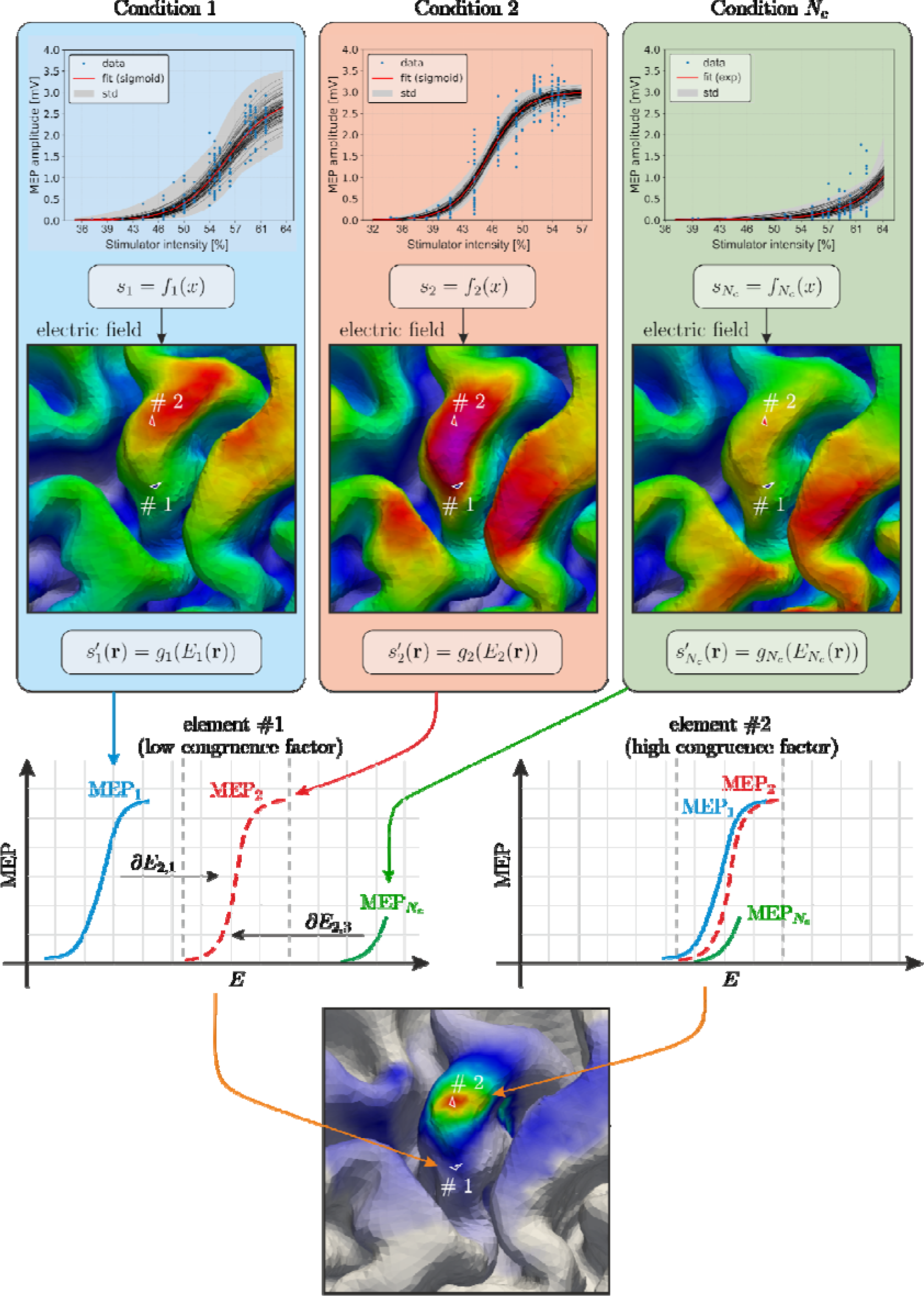
Principled approach to determine the site of stimulation by TMS. The *congruence factor* is based on the assumption that the electric field, which is causal to the observed behavioral effect corresponds for the experimental conditions. The measured I/O curves are transformed to element wis E-MEP-curves using electromagnetic field modeling (see text). The congruence factor between the E MEP-curves inversely depends on the amount of transformation (shift) necessary to obtain maximu overlap between the E-MEP-curves in each element.

The agreement between different I/O curves was quantified by computing the inverse variance of the optimal shifts τ_i_ with *i* = 1…*N*_*C*_ of the I/O curves across the experimental conditions.

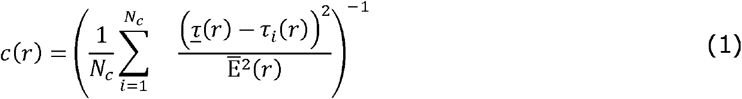

The congruence factor *c*(*r*) was calculated in each element in the ROI on the cortex by determining the inverted variance of the τ_i_, additionally weighted by the average electric field magnitude (or its normal or tangential component) squared at this location: 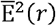. Hence, the congruence factor quantifies a relative similarity between the observations in the different experimental conditions independent on the scale of the electric field. Higher similarity between curves leads to higher inverse variance. The optimal shifts *τ*_*i*_ were obtained by determining the individual locations where the overlap against a reference curve, e.g. the first E-MEP curve, is maximized. As a result, the problem of determining the congruence factor turns into many optimization problems to calculate the shifts *τ*_*i*_ for each condition and in each element in the ROI:

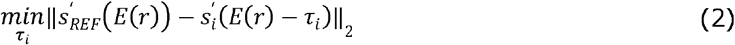

Where 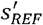 and 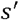 denote the reference and one of the I/O curves the shift is calculated for, respectively. This method is very general, as it is independent of the involved function types.

Because the electric field scales the I/O curves linearly, eq. (2) can be reformulated in terms of stimulator intensity, which allows a highly efficient implementation. A more detailed mathematical description is given in Section 1.2 of the *Supplementary Material*.

This is beneficial if the experimentally determined I/O curves capture only a linear or exponential part of the relationship between electric field and MEP amplitude (see above). If, however, each E-MEP curve can be represented as analytical sigmoidal function, parameterized by its turning point *x*_0,*i*_, the shifts *τ_i_* are directly given by *τ_i_* = _0,*i*_E_*i*_(*r*), and the computational expensive optimization from eq. 2 is avoided, thus making the method computationally very efficient. This approach is advantageous in terms of computational cost and is preferred in the current study if all I/O curves are modelled by sigmoidal functions.

As the standard Freesurfer average template (FsAverage) suffers from several malformed elements at the primary motor cortex with each of roughly the 20-fold size as the average elements, we created a group-based average with Freesurfer. In this iterative procedure, a randomly chosen subject was used as initial template and all other subjects were registered to this. In the second step, the template was updated based on these registrations. The third step comprised the registration of all subjects to this updated template. The second and third steps were then repeated to improve the template.

### 2.4 Uncertainty and sensitivity analysis

The congruence factor is influenced by several parameters. For example, previous studies have shown that, because of their large uncertainties, the electrical ohmic conductivities of brain tissues have a strong influence on the magnitude of the electric field (Weise et al., 2015; Codecasa et al., 2016). Furthermore, the estimated parameters of the fitted MEP curves are also uncertain due to measurement noise (Fig. 3b, grey interval). Therefore, uncertainty and sensitivity analyses are important to investigate the stability of the results and identify the parameters and their combinations with the largest impact on the results.

Since the problem is computationally complex and features a large number of parameters, an efficient approach is necessary to conduct the analysis. Here, we applied the generalized polynomial chaos (gPC) method (Ghanem et al., 2016). Its mathematical background is described in detail in Section 1.3 of the *Supplemental Material*. In short, the gPC is based on the construction of a polynomial surrogate of the congruence factor depending on the uncertain model parameters and their associated probability density functions.

Since the electric field depends on the electrical conductivities [σ] of the brain tissues, the congruence factor will be influenced by varying conductivities as well. The conductivities of GM, WM and CSF were modelled as beta distributed random variables. The impact of the other tissues, like skull and scalp on the electric field was shown to be negligible in previous studies (Weise et al., 2015; Codecasa et al., 2016; Bicalho et al., 2018). However, extending previous studies, the impact of the level of conductivity anisotropy was included in our analysis. The conductivity tensor [σ] for each voxel was derived from the diffusion tensor using the volume normalize approach (Güllmar et al., 2010). This tensor can be visualized as ellipsoid (see Fig. S2). A spherical ellipsoid represents isotropic conductivity with equal conductivity in each direction, while a cigar shaped tensor indicates that the conductivity is much larger in one direction. We implemented an anisotropy scaling factor α that transforms the diffusion tensor from the isotropic case (α = 0) via the original tensor obtained from DTI (0.5) to a very anisotropic case (1). Although, in principle, α could be different in each voxel, this would render the resulting problem intractable. Instead, we assumed that α is the same for all voxels. This reflects systematic errors and uncertainties in the transformation between the diffusion tensor, which depends on the mobility of water molecules, and the conductivity tensor, which represents the mobility of charges. A detailed mathematical description of the parametrization of the fractional anisotropy is given in *Supplemental Material*: Section 1.2.

In addition to the conductivity and anisotropy uncertainties, the turning points, from the sigmoidal I/O curves were included in the uncertainty analysis. Their uncertainties were derived from the confidence intervals of the curve fits (cf. Fig. 3b). The stochastic properties of all investigated parameters are summarized in Table 1. The model of the congruence factor used in the gPC based uncertainty and sensitivity analysis is described in detail in *Supplemental Material*: Section 1.4.

**Table 1:** Limits and shape parameters of the model parameters for subject S1

| Parameter | Description | Min | Max | p / q | Reference / source |
| --- | --- | --- | --- | --- | --- |
| $\sigma_{WM}$ | White matter (WM) | 0.1 S/m | 0.4 S/m | 3 / 3 | Li et al. (1968)<br>Nicholson (1965)<br>Akhtari (2010) |
| $\sigma_{GM}$ | Grey matter (GM) | 0.1 S/m | 0.6 S/m | 3 / 3 | Li et al. (1968)<br>Ranck (1963)<br>Logothetis et al. (2007)<br>Yedlin et al. (1974) |
| $\sigma_{CSF}$ | Cerebrospinal fluid (CSF) | 1.2 S/m | 1.8 S/m | 3 / 3 | Gabriel et al. (2009)<br>Baumann et al. (1997) |
| $\alpha$ | Anisotropy scaling | 0.4 | 0.6 | 3 / 3 | Tuch et al. 2001 |
| $x_{0,S(90^\circ)}^*$ | MEP curve | 145.9 [A/μs] | 185.9 [A/μs] | 4 / 4 | experiment |
| $x_{0,S(135^\circ)}^*$ | ——"—— | 150.0 [A/μs] | 190.0 [A/μs] | 4 / 4 | ——"—— |
| $x_{0,I(45^\circ)}$ | ——"—— | 80.7 [A/μs] | 87.9 [A/μs] | 4 / 4 | ——"—— |
| $x_{0,I(135^\circ)}$ | ——"—— | 125.7 [A/μs] | 145.7 [A/μs] | 4 / 4 | ——"—— |
| $x_{0,P(0^\circ)}$ | ——"—— | 124.2 [A/μs] | 133.3 [A/μs] | 4 / 4 | ——"—— |
| $x_{0,M1(90^\circ)}$ | ——"—— | 86.0 [A/μs] | 98.7 [A/μs] | 4 / 4 | ——"—— |

*Note: These parameters were considered in the uncertainty and sensitivity analysis to determine the site of stimulation by means of the congruence factor. The asterisk marks conditions, where only the lower tail of the I/O curve could be determined. These curves are subject to a higher measurement uncertainty.*

After deriving the polynomial surrogate using the gPC, the spatial distribution of the μ(*r*); expectation and the variance *v*(*r*)of the congruence factors *c*(*r*) can be calculated. We further analyzed the relative standard deviation 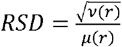 dentify possible parameter n to ranges, where the congruence factor is primarily influenced. Finally, in the sensitivity analysis, the variance was decomposed into its origins by a Sobol decomposition. The Sobol indices *S*_*i*_(*r*) represent portions of the total variance *v*(*r*), which are due to individual parameters *p*_*i*_ or a combination thereof, e.g. the conductivity of GM or the combination between different measurement parameters (Sobol, 2001; Sudret, 2008).

### 2.5 Validation

To validate the estimated sites of stimulation, we first determined for each of the three subjects of Experiment II the optimal coil position and orientation such that the cortex position at which the electric field maximum was generated coincided with the maximum of the congruence factor. This was done using an exhaustive search optimization procedure, which is described in detail the Supplemental Material 1.5 and will be implemented in the next SimNIBS major release. The identified optimal coil configurations were compared against other neighboring coil configurations by determining the MT at each site. During MT determination, single biphasic pulses with an inter stimulus interval of 5 s were applied. If the determined congruence factor hotspot is indeed the optimal site for stimulation, then the MT should lowest for the optimized coil position/orientation.

## 3 RESULTS

In the following sections, we present the results from Experiment I (15 subjects and 6 experimental conditions) and Experiment II (3 subjects and 20 experimental conditions). The latter includes a permutation analysis to determine the number of stimulation sites and corresponding coil positions required to determine the stimulation site reliably. The results from both studies are then compared. Finally, the results of the uncertainty and sensitivity analysis of the congruence factor are presented for one exemplary subject. The most influencing parameters of the numerical model and the experimental data are identified in the ensuing sensitivity analysis.

### 3.1 Experiment I (15 subjects, 6 experimental conditions)

Figure 7 shows the congruence factors of the group average and the 15 individual subjects. The electric field distributions of all conditions were determined for each subject and combined with the fitted MEP curves using the optimal curve shift approach because not all MEP curves could be fitted to sigmoidal functions. In 6/15 subjects (marked with an asterisk, *), no I/O curve could be determined for the posterior coil position P_0°_. Hence, the congruence factor was determined using only 5 of the 6 conditions. The congruence factor was calculated for the magnitude (|*E*|), as well as the normal (|*E*_┴_|) and the tangential (*E*_∥_) component of the induced electric field. The magnitude and the tangential component reached substantially higher congruence factors and smoother spatial distributions than the normal component *c*(|*E*_┴_|). In general, a clear hotspot for *c*(|*E*|) and *c*(|*E*_‖_| could be identified in the hand knob area of M1 on the gyral crown of the average template. However, considering the individual congruence factor maps shows that in 7/15 subjects (S1-S7, highlighted in Fig. 7 with a green background) we found clear and unique hotspots only on the gyral crowns in the hand knob area. In 4/15 subjects (S8-S11, Fig. 7, yellow background), we observed a second hotspot in the somatosensory cortex (S1). In 4/15 subjects (S12-S15, Fig. 7, orange background), we could only identify a dominant hotspot in S1. We reason that this is due to array ambiguities, i.e. spurious overlaps, of the realized electric fields and the missing I/O curve of condition P_0°_, in 2/4 and 3/4 subjects of the two groups, respectively. Note that maximum values of the congruence factors substantially differ across subject. This is because small differences in near zero variances among I/O curves may result in large difference in their inverse, that is, the associated congruence factors.

**Figure 7:**
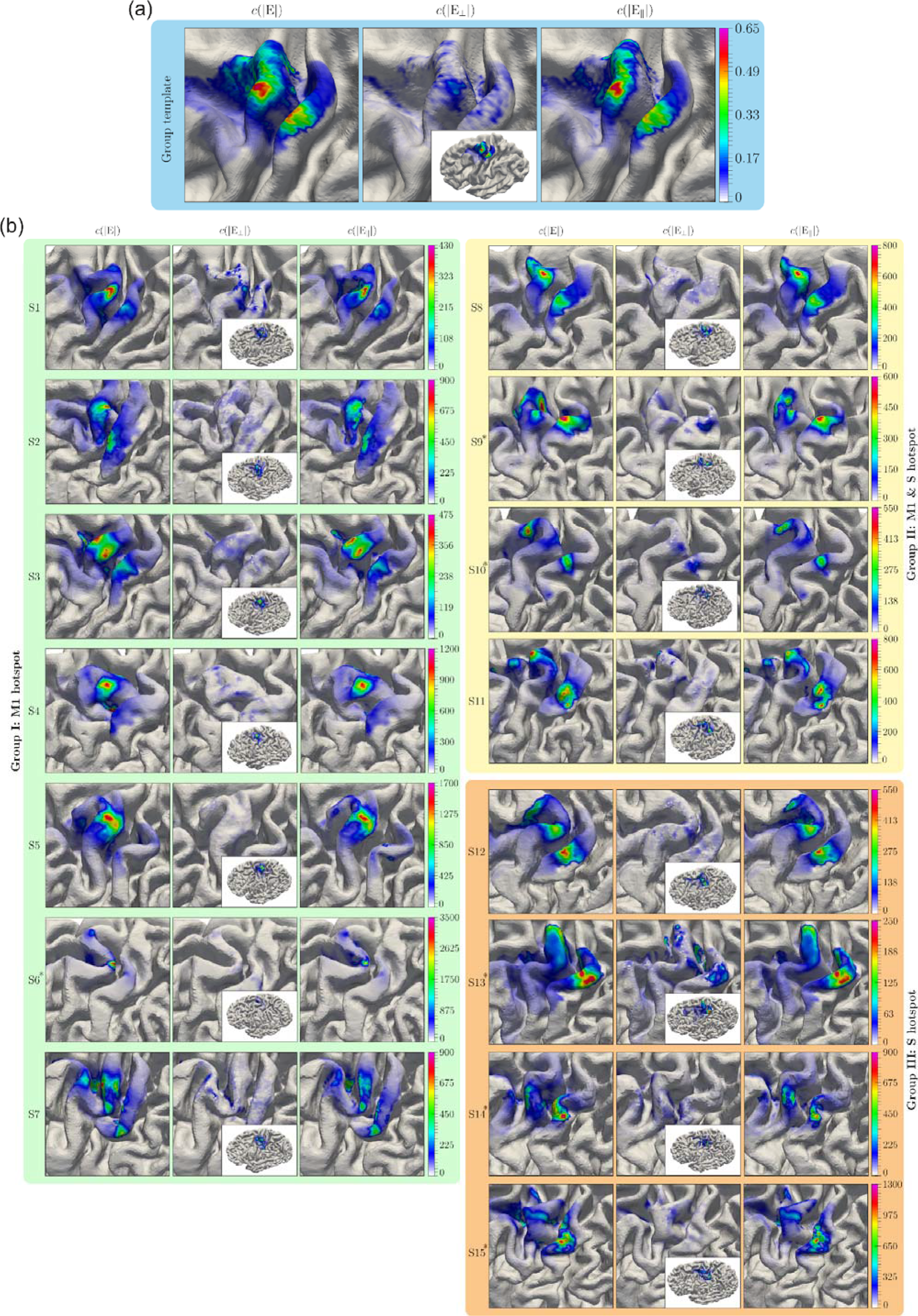
Congruence factor maps of all 15 subjects including six experimental conditions (Experiment I). The congruence factors were calculated for the magnitude, the normal and the tangential component of the electric field using the optimal curve shift approach. 7/15 subjects show unique hotspots in M1 (highlighted in green); 4/15 subjects show hotspots in M1 and S1 (highlighted in yellow) and 4/15 subjects show hotspots in S1 only (highlighted in orange); The asterisks (*) mark subjects, where no evaluable MEPs could be determined for the posterior coil position P_0°_. In these cases, the congruence factor was determined using only five conditions; all results were normalized, mapped and superimposed on the group average template shown on the top (highlighted in blue).

We expected that additional experimental conditions, i.e. more coil positions and orientations, would improve the results of the congruence factor towards more plausible hotspot locations in the M1 hand knob area. This hypothesis was investigated in Experiment II by increasing the number of coil positions and orientations from 6 (resp. 5) to 20 (cf. Fig. 2b). We selected one subject out of each of the three result groups described above (S1, S8, and S12) for this experiment.

### 3.2 Experiment II (3 subjects, 20 experimental conditions)

This experiment was conducted with an extended set of coil positions and orientations (Fig. 2b). For each subject, 20 electric field distributions were calculated and combined with the obtained MEP curves to determine the congruence factor maps. In this experiment, all I/O curves could be fitted to sigmoidal functions, which permits to avoid the computationally expensive optimizations step from (2) by directly using the variance of the turning points.

Because 20 experimental conditions are too time consuming to record in future mapping applications based on the proposed method, we investigated how the congruence factor convergences depending on the number of experimental conditions. This enabled us to determine an optimal number and selection of coil positions/orientations to reduce the experimental effort. Consequently a permutation study was performed for each subject by determining the congruence factor for all combinations of *k* = 2…20 available experimental conditions. The total number of considered conditions was 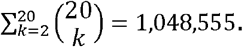. We quantified the focality of each congruence factor map by determining the area with *c* > 30. This threshold was chosen based on the permutation results data to allow comparability between the combinations and subjects. The smaller this area, the more concentrated the map is. That is, the more uniquely the causal relationship between electric field and MEP could be determined.

The results of the permutation study are shown in Fig. 8 for one exemplary subject (S1). The results for the remaining subjects were similar (see Fig. S6 and Fig. S7, respectively). We expected that a lower cross-correlation across the condition-wise electric fields would allow for a higher discriminative power in the determination of the stimulation site. This was confirmed by the analysis in Fig. 8(a), showing a correlation of *r* =0.57(*p* ≪ .001) between the size of the hotspot area and the cross-correlation of the electric fields over all *k*. As obvious from the individual number of conditions *k*, the correlation between the resulting hotspot area and the cross-correlation of the electric fields was stronger for low *k* (Fig. 8b, correlation coefficients). The median of the hotspot area converges when increasing the number of active conditions. Moreover, the spread of the area decreases by adding more information to the congruence factor calculation. Importantly, the smallest areas (lower dashed line in Fig. 8b) indicates that some condition combinations for *k*≥5 result in similar or even smaller areas than for *k* = 20. This shows that the site of stimulation can be determined with relatively few measurements by selecting optimal coil positions and orientations.

**Figure 8:**
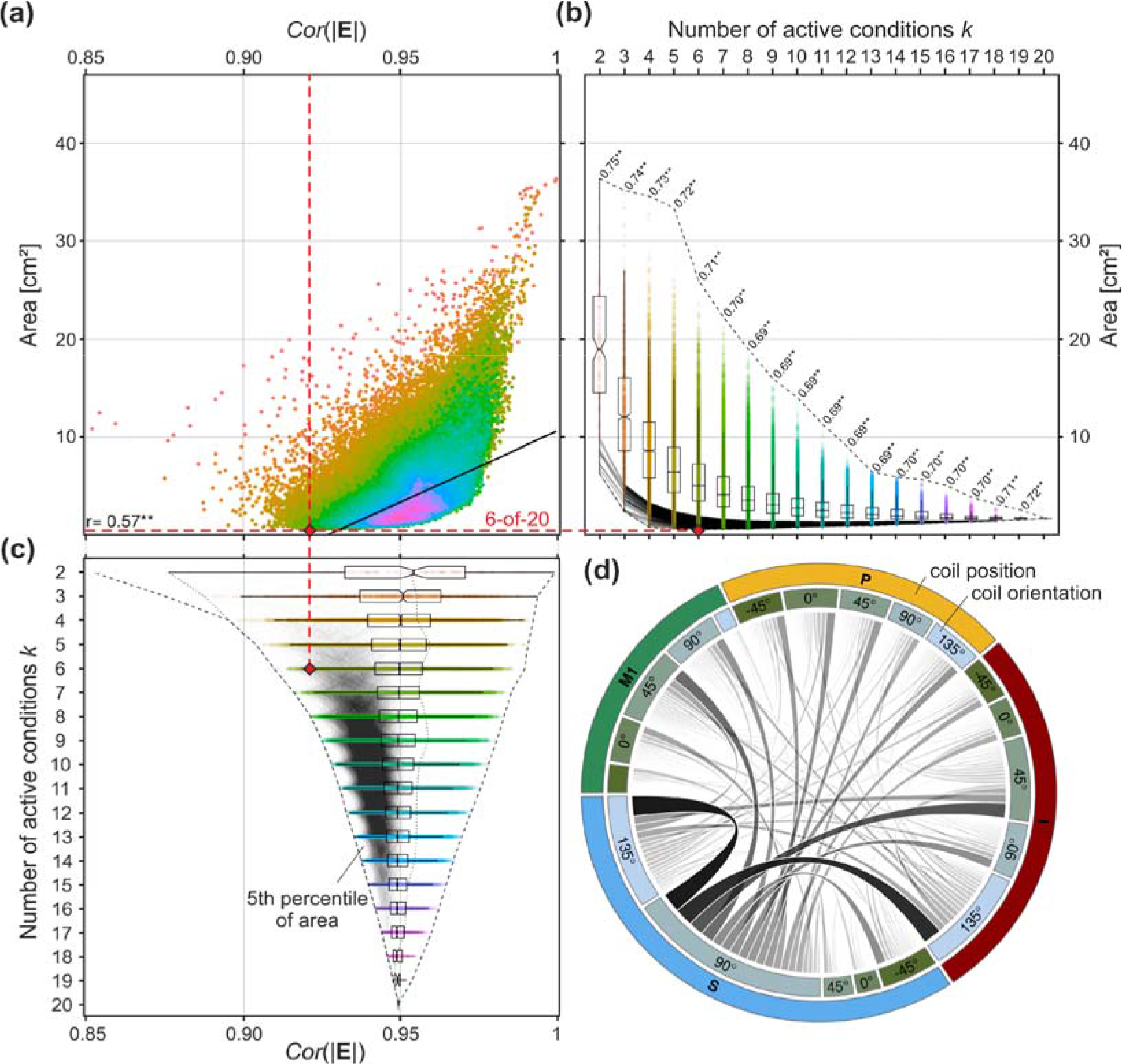
Permutation analysis interrelating the cross-correlation of the electrical fields from the different experimental conditions of Experiment II for subject S1 with the corresponding hotspot area. The hotspot area was defined as the region where *c* > 30. For each case *k*, the congruence factor was determined (20*k*) times. (a) Relationship between the cross-correlation of the electric field magnitude and the resulting hotspot area size. Colors: active conditions *k*. The correlation coefficient between the hotspot size and the cross-correlation of the electric fields over all *k* (black line) is *r* × 0.57(p ≪ .001). (b) Boxplot pf the hotsspot and area of the congruence factor depending on the number of active conditions. Box areas indicate the 25% to 75% quantiles with notch at median. Correlation coefficients between the hotspot size and the cross-correlation of the electric fields for each *k* are given. ** depict *p* < .01 (after Bonferroni correction). Grey lines: 5^th^ percentile of the best condition combinations for each *k*. Dashed lines: absolute range. The variation of the hotspot area size decreases with increasing. (c) Relationship between the cross-correlation coefficient of the electric fields the number of active conditions *k*. Box areas indicate the 25% to 75% quantiles with notch at median. Grey lines: 5^th^ percentile of the best condition combinations for each *k*. Dashed lines: absolute range. The dashed red lines highlight the case of the 6 coil positions and orientations, where the congruence factor map was most focal. Its cross-correlation coefficient is with 0.921 lower than the first quartile of possible solutions. (d) Chord graph highlighting the interaction and relative contribution between different coil positions (outer circle) and coil orientations (inner circle) from the 5^th^ percentile of best condition combinations over all *k* resulting in small hotspot areas (highlighted with black lines in (b)).

The relationship between the cross-correlation of the electric fields and the number of active conditions *k* is shown in Fig. 8c. Cases resulting in the smallest 5^th^ percentile of the hotspot area are shown as thin lines in the shaded area, corresponding to the ones in Fig. 8b. As expected, these cases are concentrated in regions of low cross-correlation.

The data were further analyzed to identify which combinations of experimental conditions were especially informative and produce very focal hotspots (Fig. 8d). This analysis was performed for *k* = 6. The appearance of each condition and its co-occurrence with other conditions was accumulated across all condition combinations, which are part of the smallest 5^th^ percentile of the hotspot area (grey shaded area in Fig. 8b). We observed that the co-occurrence was not random and combinations surrounding M1, i.e. inferior, superior, and posterior, appeared more often than coil positions directly over M1. The corresponding coil orientations considerably differed and connections between S_90°_, S_135°_, I_45°_, and I_135°_ stand out. Moreover, it can be observed that posterior conditions occurred frequently in combination with S_90°_, which further confirms the need for highly varying electric field distributions. This behavior is even stronger pronounced for subjects S8 and S12 (cf. Fig. S6 and Fig. S7 in the *Supplemental Material*).

In the following, the results for *k* = 6 condition combinations out of the 20 experiments (6-of-20) resulting in the smallest hotspot area are described in more detail and compared to Experiment I and to the full 20-of-20 result. The results are shown in Fig. 9 for each subject. The congruence factor maps were normalized with respect to their individual maxima to allow comparability. For subject S1, we already found a unique hotspot in the M1 hand knob area in Experiment I. The results of Experiment II show that this pattern is reproducible and even more focused (as the deflection on the somatosensory cortex is weaker) for the best combination of 6-of-20 conditions. Hence, for this subject, the coil positions of Experiment I were already sufficient to determine the site of stimulation in a plausible manner. The second subject belongs to the group, which showed hotspots in both M1 and S1 (Experiment I). In Experiment II, a single hotspot was limited to the M1 region as well, and the deflection in S1 disappeared. The M1 hotspot was also slightly shifted inferior. The third subject belongs to the group which showed a hotspot only in S1 in Experiment I. In Experiment II, however, the hotspot moved to M1 supporting our assumption of insufficient information content concerning the combination of electric field profiles and measured MEP amplitude curves due to a limited classification ability of the electric fields. As indicated by the convergence results of the permutation study (Fig. 8a), adding the remaining conditions of Experiment II (20-of-20 case) does not yield any improvement for any of the three subject groups.

**Figure 9.**
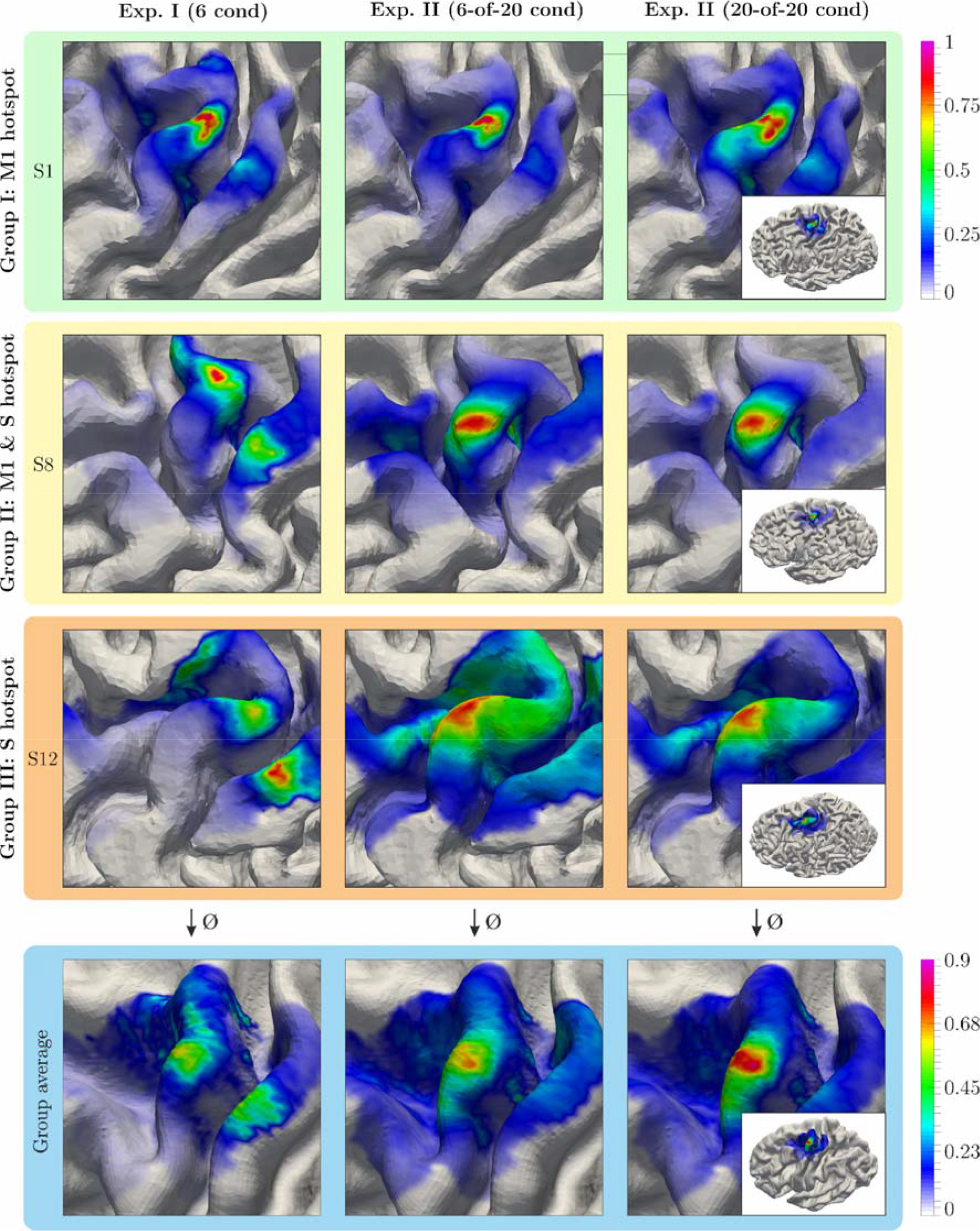
Normalized congruence factor maps of 3 subjects. The first column shows the results of Experiment I with 6 experimental conditions; the middle row depicts the 6-of-20 condition combination with the smallest hotspot area from Experiment II; The right column shows the congruence factor maps when all 20 experimental conditions of study II are included in the analysis. The subjects were chosen to include on participant from each result group in study I (M1 only, M1 and S1, S1 only, see Fig.7).

For subject S1 (first row in Fig. 9), the cross-correlation of the electric field distributions in the ROI was 0.951 for Experiment I, and 0.921 for Experiment II. For subject S8 (second row in Fig. 9), the cross-correlation was 0.953 and 0.925 for Experiment I and II, respectively. We observed that the use of less correlating electric field distributions increased the quality of the reconstruction. Finally, for subject S12 (third row in Fig. 9) the cross-correlations were nearly the same with 0.951 and 0.953 from Experiment I to II, respectively. However, the improvement of the results indicates that the selected coil positions and orientations in Experiment II were more suitable to determine the congruence factor, resulting in a higher distinguishability between the cortical positions in Experiment I. We wish to emphasize that this property is only partly reflected by the cross-correlation coefficient. A definition of a more sophisticated ambiguity measure to determine an optimal set of coil positions and orientations will be subject of a future study.

### 3.3 Validation

After determining the optimal coil positions and orientations for the subjects in Experiment II, we validated the predicted cortical sites of stimulation. For health reasons not related to this study, subject 12 from group III (c.f. Fig. 9) was not able to participate in the validation study. We replaced that subject by subject 15 from group III and repeated Experiment II. It turned out that, in this subject, using the predefined 20 conditions did not yield a single pronounced congruence factor hotspot. To increase the electric field variance, we added further conditions at different positions, orientations, and tilting angles of the TMS coil (see Fig. S5 of *Supplemental Material*). We additionally determined the corresponding best 6-of-30 condition combination yielding very similar results compared to the result to the full 30 condition analysis. The results are shown in Fig. 10.

**Figure 10.**
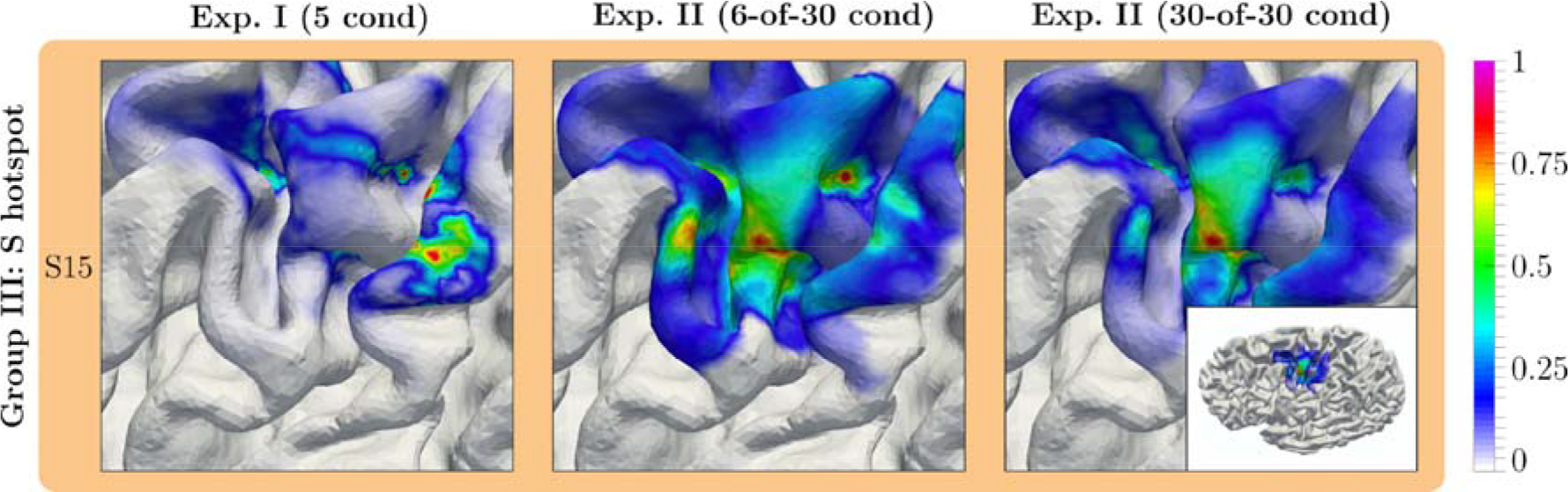
Normalized congruence factor maps of subject 15, replacing subject 12 from subject group III for the validation. The first column shows the results of Experiment I with 5 experimental conditions; the middle row depicts the 6-of-30 condition combination with the smallest hotspot area from Experiment II; the right column shows the congruence factor map when all 30 experimental conditions of study II are taken into account.

We measured the lowest MTs at these optimal coil positions and orientations compared to all other tested coil configurations (Fig. 11). Notably, all computationally determined optimal coil orientations are fairly similar to the commonly used 45° coil orientation towards the *fissura longitudinalis* (Brasil-Neto et al., 1992; Mills et al., 1992).

**Figure 11.**
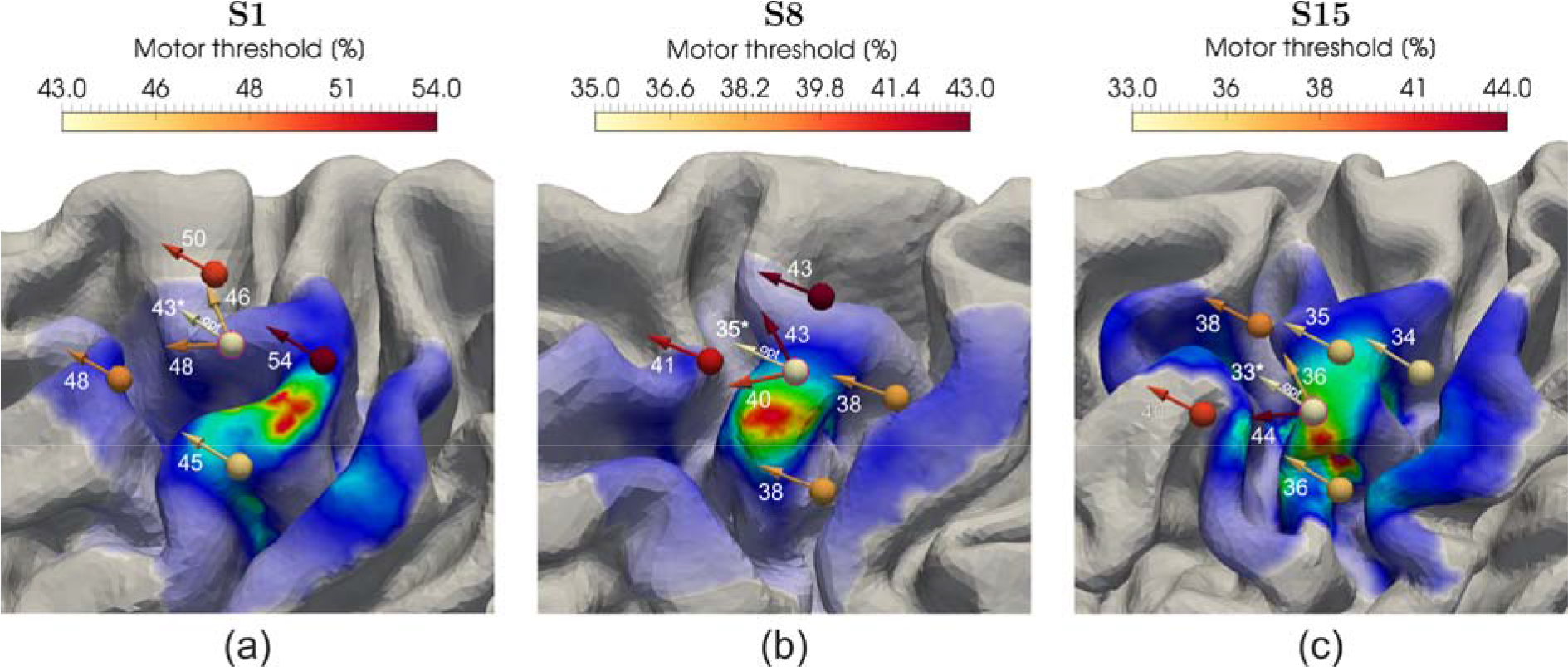
Coil positions and orientations used to validate the determined cortical site of stimulation. The optimal coil position is marked with a dashed purple circle and its corresponding optimal orientation is indicated by “opt”. Numbers represent the resting motor thresholds determined where 5 of 10 consecutive MEPs reached values > 50 µV. Lowest MTs at the optimal coil positions and orientations are marked with an asterisk.

### 3.4 Uncertainty and sensitivity analysis

We analyzed the congruence factor results in terms of uncertainties and sensitivities towards the electrical conductivities of brain tissues, fractional anisotropy, and measurement inaccuracies for all subjects of Experiment II considering the 6-of-20 case described previously. The results from subject S1 are shown in Fig. 11. The results of subject S8 and S12 are shown in Fig. S8 and Fig. S9. The uncertainties of the model parameters are listed in Table 1 and Table S1 & S2. The spatial distributions of the mean, the relative standard deviation (*RSD*), and the variance (*VAR*) of the congruence factor are shown in Fig. 12a. The mean distribution shows a hotspot, which is extending from the gyral crown of M1 to upper parts of the anterior sulcal wall. *RSD* and *VAR* indicate that, the congruence factor could be determined with a greater certainty (*RSD* ≈ 12%) on the gyral crown than on the anterior sulcal wall (*RSD* ≈ 40%).

**Figure 12:**
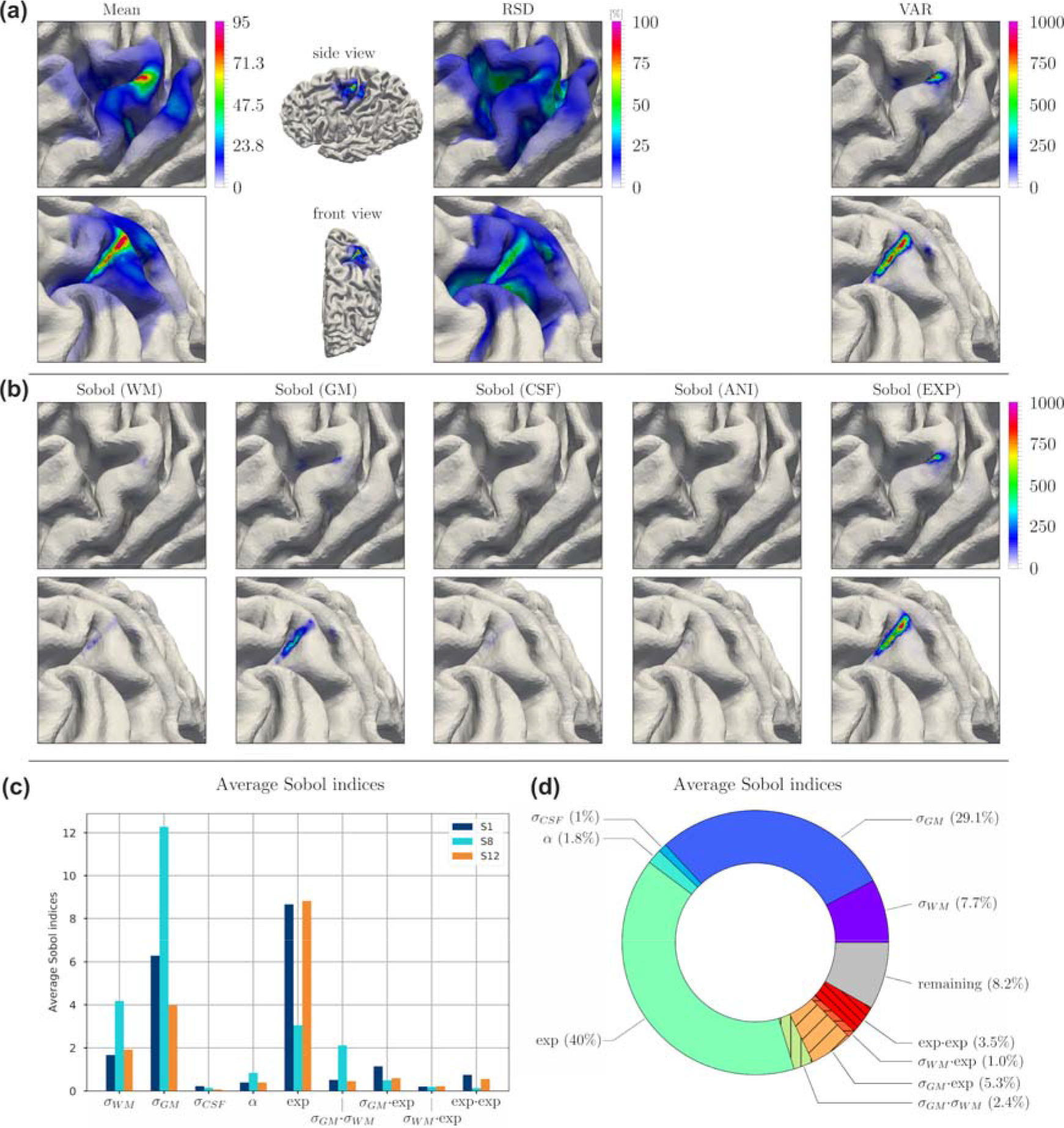
Uncertainty and sensitivity analysis of the congruence factor determined for subject S1 in the 6-of-20 analysis of Experiment II. (a) Expected value (Mean), relative standard deviation (RSD), and variance (VAR) of the congruence factor. (b) Spatial distributions of the absolute first order Sobol indices. The normalization with respect to the total variance was avoided to strengthen the focus on regions of high variance (see eq. (12) in the *Supplementary Material* on how the Sobol indices are computed). The Sobol index maps of the individual MEP parameters resulting from uncertainties in the experimental data are summarized into one Sobol index map “Sobol (EXP)”. (c) Average first order Sobol indices for subjects S1 (shown here), S8, and S12. The average was computed over the elements in the ROI (d) Relative first order Sobol indices averaged over the ROI. For (a) and (b) two different perspectives are shown (top and bottoms rows), in order to improve visibility of the effects.

To identify the most influential model parameters, we decomposed *VAR* into its origins by a Sobol decomposition. The spatial distributions of the absolute first order Sobol indices on the cortex are shown in Fig. 12b. The Sobol coefficient maps of the MEP curve parameters are accumulated to one Sobol index termed Sobol (EXP). The average first and highest second order Sobol indices are depicted in Fig. 12c for all subjects of Experiment II. The average was computed over the elements in the ROI. The most protrude parameters that contribute to the uncertainty of the congruence factor were the electrical conductivities of GM (σ_*GM*_), WM (σ_*GM*_) as well as the uncertainty of the measured MEP curves (EXP) for all subjects. Their relative contribution to the total variance, shown in Fig. 12 d (and in Fig. S8d and S9d), was subject specific and depended on the accuracy of the measured MEP curves as well as the brain anatomy, influencing the electric field distribution. The uncertainty of the congruence factor hotspot in the anterior sulcal wall predominantly originated from the uncertainty of σ_*GM*_ and the measurement uncertainties indicate that this hotspot is likely to be spurious.

## 4 DISCUSSION

### 4.1 Summary of Findings

In the present study, we introduce a novel approach that links numerical modeling of the induced electric field with measurements of peripheral physiological responses to considerably improve the localization of effectively stimulated areas during TMS. With this approach, we were able to accurately localize the cortical area that is responsible for the observed motor output when single TMS pulses were applied over the motor cortex. Our main finding was that sharply bounded neural structures located in the gyral crowns extending to the upper parts of the sulcal wall of the motor hand area represent the most likely origin of the motor evoked potentials. We identified the magnitude and the tangential component of the electric field as the relevant quantities for modulating the observed effect.

Our results implicate that unique results can be obtained with relatively few measurements. Based on our findings, we derive principles for the selection of the respective coil positions that may help to improve localization of TMS effects in future applications, both at the single subject and group level. Our first experiment combined two different stimulation sites with three coil rotations each, yielding 6 conditions. The induced electric fields were computed with FEM, allowing for the assessment of element-wise E-MEP relations. We identified three groups with a hotspot at either M1, the somatosensory cortex, or both. One subject from each group was included in the extended second experiment. For the subject from the M1-group, the hotspot was replicated at the same spot. For the subjects from the other two groups, the results could be significantly improved and single hotspots at the gyral crowns and the upper parts of the sulcal walls of the motor hand area were observed in all cases. The final validation study in three subjects confirmed that optimizing the TMS coil position and orientation such that it maximized the electric field at the predicted cortical target indeed resulted in a minimization of the MTs.

The congruence factor employed in our study quantifies the correlation between the measured physiological variable (here, the MEP) and the induced electric field profiles. Note that the proposed approach does not depend on the involved function types to describe the I/O behavior. This provides a high level of flexibility and makes the method easy to adapt to other applications and domains. We conclude that our approach significantly improves the localization of effectively stimulated areas during TMS and may increase the power and reliability of the resulting effects in future TMS studies at the individual level.

### 4.2 Relation to prior studies relating TMS electric fields with MEPs

Based on our results, we argue that areas with maximum congruence factors are good candidates for effective stimulation. Importantly, in all subjects, we observed that higher variability between electric fields sharpened the localization results. Sets of experimental conditions that selectively varied coil position or coil orientation did not contain sufficient information to uniquely determine the effective cortical stimulation site. Moreover, stimulation directly over M1 with 45° orientation, though yielding the strongest effect, was by far not the most informative condition in our method, which can be explained by the relative wide spread of the electric field in the motor area, and hence, with low discriminative power, produced by standard figure-of-eight coils. These observations might provide a potential explanation for the spurious second hotspot in the somatosensory cortex observed by Bungert et al. (2017) when stimulating selectively over M1 and Laakso et al. (2018), stimulating with a 45° coil orientation towards the *fissura longitudinalis* (Brasil-Neto et al., 1992; Mills et al., 1992). Notably, we observed similar effects in Experiment I, where only 5 or 6 non-optimal experimental conditions were considered (subject group II and III, Fig. 7). Reducing the correlation of the electric fields across the tested positions and orientations considerably enhanced the localization capabilities of our method in all subjects (Experiment II). This observation was further supported by a permutation analysis showing that higher variability between the spatial patterns of the electric fields, by means of using particular combinations of coil positions and orientations, considerably increased the accuracy of the localization results.

Interestingly, studies incorporating selectively the 45° coil orientation towards the fissura longitudinalis (Laakso et al., 2018; Krieg et al., 2013; Salinas et al., 2011) appear to support sulcal wall activation by the normal component of the electric field. In contrast, studies which involve different coil orientations (Bungert et al., 2017) highlight |*E*| and gyral crowns. Recent results from direct electric stimulation (Aonuma et al., 2018) support the notion of gyral crown activation, which contrasts with the conclusions drawn from applying imaging techniques (Fox et al., 2004; Krieg et al., 2013). However, both methods have the major disadvantage that their resolution in the current state of research is not sufficient to answer this question. By changing both, i.e. coil position and orientation, we observed low congruence factors for the normal component of the electric field at the anterior wall of the central sulcus. Since low congruence factors highlight areas where the behavioral effect does not correlate with changes in the local electric field, our results indicate that the previously proposed stimulation mechanism by the normal component (Laakso et al., 2018; Fox et al., 2004; Krieg et al., 2013) cannot explain the observed effect for all experimental conditions. In contrast, we found that the tangential component (and therefore also the magnitude) of the field showed reasonable congruence factor maps. This finding suggests that the gyral crowns and upper parts of the sulcal wall are the most likely origin of the motor evoked potentials.

Two prior studies superimposed the calculated electric fields either in additive or multiplicative fashion to localize the cortical position targeted by TMS (Opitz et al., 2013; Aonuma et al., 2018). Opitz et al. (2013) weighted the computed electric fields with the strengths of the observed effects and overlaid the fields in an additive fashion. In contrast, Aonuma et al. (2018) superimposed the fields in a multiplicative fashion after selecting the experimental conditions for which the observable effect exceeded a particular threshold. The latter may be disadvantageous since it uses only a small portion of the information contained in the measurement. Both methods approximate a covariance between the field strength and the MEP amplitude. However, this covariance does not only depend on the correlative relationship between the two, but also on the general magnitude of the field across conditions. This leads to a strong bias towards voxels which generally receive higher field strengths (i.e., on gyral crowns) for both approaches.

Our validation study confirmed the general optimality of the PA-45 coil orientation towards the *fissura longitudinalis* (Brasil-Neto et al., 1992; Mills et al., 1992). The slight deviations between the optima confirm the inter-individual variability in optimal coil orientation observed for example by Balslev et al. (2007) and Bungert et al. (2017).

### 4.3 Relation to simulations of neural excitation by TMS

Combining electric field computations and compartment models of neurons, Seo et al. (2017) propose the initial segments of pyramidal cells in layer 3 and 5 to be the sites of effective stimulation, possibly due to the omnidirectional orientation of the basal dendritic trees. Alternatively, also the terminals of axon collaterals might be stimulated (Aberra et al., 2018), which again are equally distributed in all directions around the main axon and have low thresholds. Our results, namely high congruence factors of the magnitude and the tangential component of the electric field in the gyral crown and rim, indicate that the stimulation mechanism of TMS may indeed occur due to synaptic or dendritic activation of neurons. This is in line with predictions from previous modeling studies (Silva et al., 2008; Salvador et al., 2011). Future studies may extend the congruence factor approach to more detailed neuron models (Moezzi et al., 2018) and tractography-based fiber tracts (De Geeter et al., 2015; De Geeter et al., 2016).

### 4.4 Factors influencing the stability of the results

The uncertainty and sensitivity analyses confirmed robust hotspots on the gyral crowns extending to upper parts of the sulcal wall of M1 (cf. Fig. 10, Fig. S8, and Fig. S9). The maxima of the means coincide well with the results of the deterministic case (cf. Fig. 9). The relative standard deviation (RSD) in the hotspots on the gyral crowns varies between 10-25%, depending on the subject. The uncertainties mainly translate into uncertainties of the congruence factor on the anterior sulcal wall of the precentral gyrus and not on the gyral crown, where the primary hotspot was detected.

The hotspots on the anterior sulcal wall are lying in line with the normal vector from the head surface towards the center of the brain. We hypothesize that these spurious hotspots are projections from the gyral crown hotspots. Since our approach is independent of the magnitudes of the electric field, but sensitive to their spatial profiles, these spurious hotspots might result from insufficient electric field variance between these locations.

Decomposing the variance into their origins revealed a strong contribution from GM and WM conductivity as well as from the measured I/O curves. This is in line with previous studies, which showed that the electrical conductivities of GM and WM are the most influencing parameters considering the induced electric field in grey matter (Weise et al., 2015; Codecasa et al., 2016; Saturnino et al., 2018). The impact of the measurement uncertainty was lower for subject S8 compared to the other subjects, which can be explained by the fact that the MEP curves could be determined with a higher certainty (cf. Table 1, S1, and S2). Nevertheless, its contribution was still high and special care should be taken when recording characteristic regions of the I/O curve, like the turning points of the sigmoids, to reduce its influence on the congruence factor. In contrast, the conductivity of CSF and the level of anisotropy had a small impact on the congruence factor and could be treated as deterministic in future analyses.

### 4.5 Towards a clinically suitable, principled TMS mapping procedure

To enable clinical applicability of the proposed method, for instance in presurgical mapping, the experimental effort has to be reduced to a minimum while ensuring reliability. Regarding our second research question, the permutation analysis from Experiment II (Fig. 8, Fig. S6, and Fig. S7) revealed that six stimulation conditions at three different locations around M1 with different orientations are sufficient to address the localization problem at hand. Notably, the actual condition combinations that result in a minimum hotspot area differ strongly between subjects. This is likely due to inter-individual differences in anatomy and functional brain organization. Using a high number of experiments increases the stability and reliability of the solution. However, at the same time, it also reduces the resolution by introducing more measurement uncertainty. This became evident in the permutation study in Fig. 8b, where the minimal hotspot area had a minimum for *k* = 6 conditions and slightly increased for higher values of *k*. Increasing the field variability is a promising starting point for subject specific optimization to determine the optimal number and selection of coil positions and orientations before the experiment. An even more sophisticated scheme could involve maximizing the distinguishability between voxels based on their sensitivity profiles, that is, the vectors of E fields caused by the different coil positions and orientations with identical activation strength. In order to distinguish two voxels with respect to their congruence factor, their sensitivity profiles should be as different as possible. The formulation of an optimization procedure that identifies the best combination of coil positions and orientations to maximize the differences of the sensitivity profiles between any two voxels in the region of interest will be investigated in detail in future research.

Beyond the localization of the origin of MEPs, our approach allows to localize functionally involved cortical areas for other processes, provided that it is possible to observe a quantitative response variable that depends on the stimulation intensity. Considering adapted experimental paradigms, which are able to capture this, future studies may use our approach for pre-surgical language or somatosensory mapping purposes.

### 4.6 Study Limitations

Our results indicate that possible carry-over effects of stimulation in Experiment II due to the relatively short inter stimulus interval (ISI) of 4 s do not affect our conclusions, as the ISI was kept stable during a single experiment and any carry-over effect should be stable as well. Therefore, the correlative relationship between electric field and MEP amplitude should remain unaffected, even if the absolute value of the MEP is changed.

So far, our method relies on the assumption that the experimental effects can be explained by activity in a single cortical patch. This holds in the current motor experiment identifying the cortical origin of FDI activation. In other experimental paradigms, however, several network nodes may exist that might influence the effect. These nodes may also influence each other in different ways, which would lead to partial correlations. Incorporating connections into our model will tremendously increase the computational cost and efficient algorithms have to be developed to combine the electric field profiles and the physiological response data. Since numerous connections can be analyzed independently from each other, the problem is highly parallelizable and well suited for GPU or HPC implementations. The extension of our technique to identify multivariate relationships between externally observable effects and stimulation of neural populations is subject of ongoing work.

## Acknowledgments

Konstantin Weise; Max Planck Institute for Human Cognitive and Brain Sciences, Stephanstr. 1a, 04103 Leipzig, Germany; Technische Universität Ilmenau, Advanced Electromagnetics Group, Helmholtzplatz 2, 98693 Ilmenau, Germany, phone: +49 341 9940-2580. This work was partially supported by the German Science Foundation (DFG) (grant number WE 59851/1); Lundbeckfonden (grant no. R118-A11308), the NVIDIA Corporation (donation of two Titan Xp graphics cards to GH and KW) and NovoNordisk fonden (grant no. NNF14OC0011413).

