## supplemental material for "A novel approach to localize cortical TMS effects"

**A novel approach to localize cortical TMS effects
Supplemental Material**

Konstantin Weise^1,2+*^, Ole Numssen^3+^, Axel Thielscher^4,5^, Gesa Hartwigsen^3#^, Thomas R. Knösche^1,6#^

*^1^Max Planck Institute for Human Cognitive and Brain Sciences, Stephanstr. 1a, 04103 Leipzig, Germany.*

*^2^Technische Universität Ilmenau, Advanced Electromagnetics Group, Helmholtzplatz 2, 98693 Ilmenau, Germany.*

*^3^Lise Meitner Research Group “Cognition and Plasticity”, Max Planck Institute for Human Cognitive and Brain Sciences, Stephanstr. 1a, 04103 Leipzig, Germany.*

*^4^Danish Research Centre for Magnetic Resonance, Centre for Functional and Diagnostic Imaging and Research, Copenhagen University Hospital Hvidovre, Denmark.*

*^5^Technical University of Denmark, Center for Magnetic Resonance, Department of Health Technology, Kongens Lyngby, Denmark.*

*^6^Technische Universität Ilmenau, Institute of Biomedical Engineering and Informatics, Gustav-Kirchhoff-Straße 2, 98693 Ilmenau, Germany.*

*^+,#^ contributed equally*

* CORRESPONDING AUTHOR

*Konstantin Weise; Max Planck Institute for Human Cognitive and Brain Sciences, Stephanstr. 1a, 04103 Leipzig, Germany; Technische Universität Ilmenau, Advanced Electromagnetics Group, Helmholtzplatz 2, 98693 Ilmenau, Germany;, phone: +49 341 9940-2580*

**1 METHODS**

1.1 **Determining the site of stimulation**

In the following, the two approaches to determine the congruence factor are presented. The site of effective stimulation is determined by combining the measured I/O-curves of the MEP amplitude with the simulation results of the induced electric field. An operator $O$ is derived to transform the brain-wide MEP amplitude curves $s_{i}=f_{i}\left( x \right)$ with $i=1,\ldots,N_{c}$ indicating the experimental condition, given as a function of stimulator intensities *x,* to an element-wise quantity $c\left( \boldsymbol{r} \right)$, given in the electric field space, i.e. $c\left( \boldsymbol{r} \right)=O\left( s\left( x \right) \right)$. The quantity $c\left( \boldsymbol{r} \right)$ is termed the *congruence factor*. The basic principle to determine $c\left( \boldsymbol{r} \right)$ is shown in Fig. 5. For each experimental condition, the corresponding electric field distribution is determined. Due to the linear relationship between the stimulator intensity and the magnitude of the induced electric field, the MEP curves can be projected and accordingly scaled to each element in the cortex. As a result, the global stimulator intensity vs MEP curves are transformed from the stimulator intensity space $s_{i}=f_{i}\left( x \right)$ to the electric field space $s_{i}^{'}=g_{i}\left( E_{i}\left( \boldsymbol{r} \right) \right)$, containing spatial information. Therefore, a series of MEP curves is assigned to each element. The definition of the congruence factor is based on the assumption that the causative electric field, which stimulates the neuronal population of interest resulting in the observed behavioral effect is stable across the experimental conditions. For this reason, we are seeking the cortical region, where all MEP curves are similar and assume that this region corresponds to the site of stimulation. This principle is illustrated in Fig. 6, highlighting an element with a low congruence factor, where the E-MEP curves are strongly diverging from each other and an element with a high congruence factor, where the E-MEP curves are similar. The congruence factor is computed in the ROI considering the magnitude of electric field $|\mathbf{E}|$ as well as its normal (${|\mathbf{E}}_{\perp}|$) and tangential (${|\mathbf{E}}_{\parallel}|$) components. Two different approaches can be applied to determine the congruence factor, which are described in the following.

**1.2 Efficient implementation of the general curve shift approach to determine the congruence factor**

Originally, the I/O curves are given in terms of stimulator intensity. The congruence factor however, has to be determined with respect to electric field. This is done by transforming the measured MSO-MEP curve for each condition (I/O curve in %MSO, and MEP amplitude) to E-MEP-curves representing the relationship between the electric field at a particular cortex location and the respective MEP amplitude.

The congruence factor is calculated by determining the shifts $\tau_{i}$ between each I/O curve and in each element:

$\underset{\tau_{i}}{argmin}\left\| s_{REF}^{'}\left( E\left( \boldsymbol{r} \right) \right)-s_{i}^{'}\left( E\left( \boldsymbol{r} \right)-\tau_{i} \right) \right\|_{2}$	(1)

Where $s_{REF}^{'}$ and $s^{'}$ denote the reference and one of the I/O curves the shift is calculated for, respectively. This method is very general, as it is independent of the involved function types.

The linear scaling of the stimulator intensity axis with the electric fields in the elements affects the shape of the I/O curves. When considering an element with a very low electric field as example, the respective I/O curve is located in the lower range of the electric field axis and stays narrow even if the intensity increases. In contrast, in an element with a very high electric field, the I/O curve is located in the high electric field range. When increasing the stimulator intensity, the E-MEP curve broadens compared to the former case. We term the effect of narrowing and broadening the *stretch effect*. This has to be accounted for when determining the curve shifts. The procedure to calculate the stretch-corrected total shift $\tau$ is illustrated in Fig. S1.

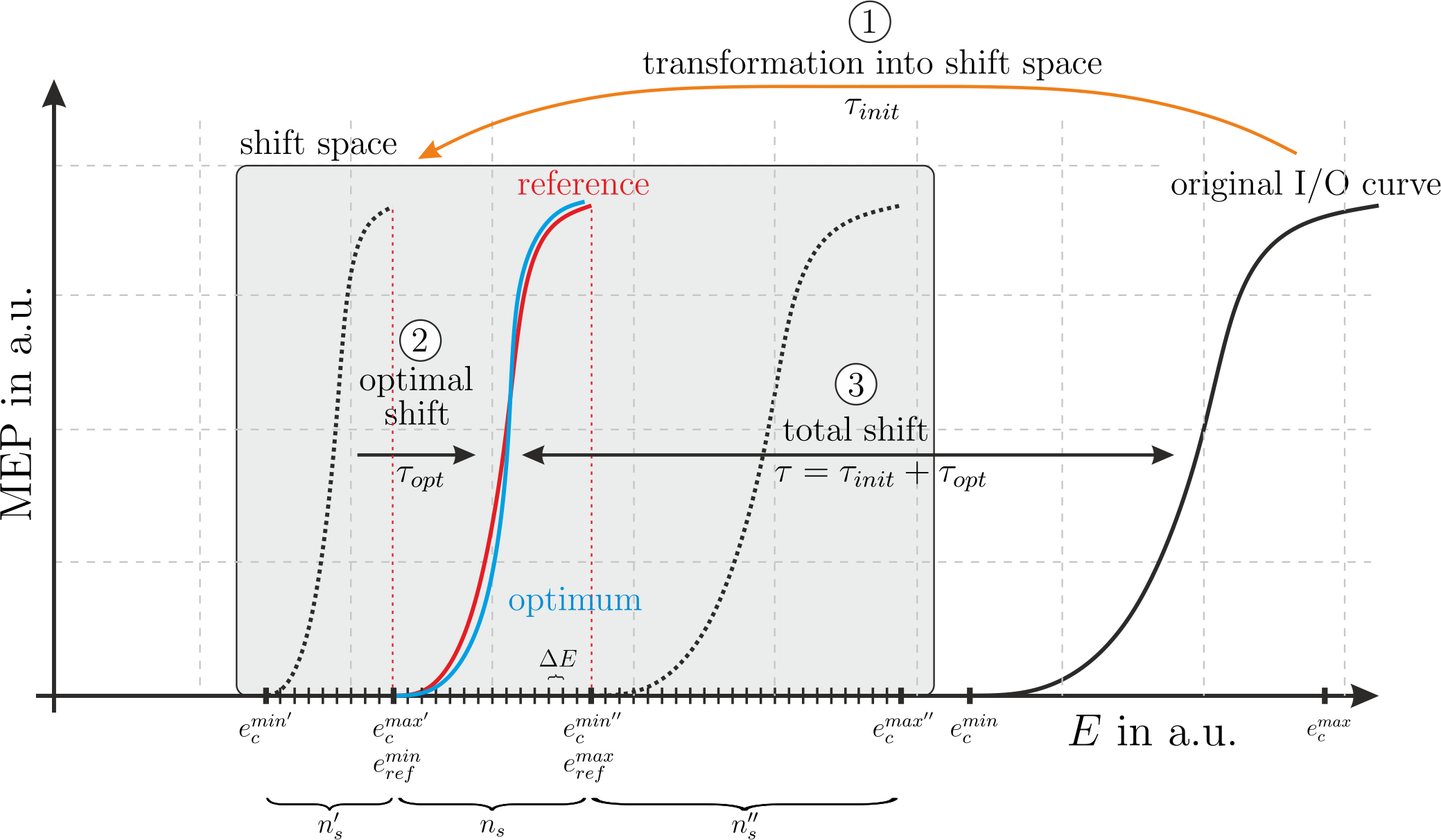

**Figure S1:** Procedure to determine the curve shifts in the congruence factor calculations. Two I/O curves are exemplary shown in the electric field space for one element in the cortex. The total shift $\tau$ between the reference (solid red) and an I/O curve under investigation (solid black) is determined as follows: (1) the I/O curve is transformed into the shift space by performing an initial shift $\tau_{init}$ including a stretch correction; (2) the transformed curve is stepwise shifted while stretch-corrected to determine the shift $\tau_{opt}$ where the curves are overlapping optimally in a mean square sense (solid blue); (3) the total shift $\tau$ is determined by adding the initial shift to the optimal shift distance. $\Delta E$ denotes the step size and is determined by the number of steps $n_{s}$ of the reference curve; $e_{c}^{min}$ and $e_{c}^{max}$ denote the minimal and maximal values of the original E-MEP curve; $e_{c}^{min'}$ and $e_{c}^{max'}$ denote the minimal and maximal values of the transformed E-MEP curve in the shift space at the starting point before the optimal displacement is determined, $e_{c}^{min''}$ and $e_{c}^{max''}$ denote the minimal and maximal values of the E-MEP curve in the shift space at the end point of the reference curve; $e_{ref}^{min}=e_{c}^{max}$ and $e_{ref}^{max}=e_{c}^{min''}$ are the minimal and maximal values of the reference curve; $n_{s}^{'}$ and $n_{s}^{''}$ denote the number of steps before and after the reference curve, respectively.

The method to determine the curve shifts $\tau_{i}$ consists of the following steps:

I. Transform the I/O curves from the stimulator intensity space to the electric field space by weighting each abscissa with the corresponding element- and condition wise electric field.
II. Define a reference curve to define the shift space (e.g. the first I/O curve).
III. Transformation of the remaining I/O curves into the shift space (determine the initial shifts and stretch).
IV. Successively shift the I/O curves with respect to the reference curve and determine the shifts where the curves are overlapping optimally.
V. Determine the total shifts by adding the initial shifts to the optimal shifts.

The initial shift $\tau_{init}$ (including the stretch correction) depends on the electric fields and has to be calculated for every condition (excluding the reference) and in every point on the cortex. It can be efficiently expressed in matrix notation by:

${[\tau}_{init}]=\left[ E_{c}^{1\%} \right]\circ\left[ I_{c}^{min} \right]-I_{ref}^{min}\mathbf{e}_{ref}^{1\%}\otimes{(\mathbf{I}_{c}^{min}⊘\mathbf{I}_{c}^{max})}^{T}$ ,	(12)

where ${[\tau}_{init}]$ is a matrix containing the initial shifts of all $N_{cond}-1$ remaining E-MEP curves in every ROI element and is of size $[N_{ROI}\times(N_{cond}-1)]$; $\left[ E_{c}^{1\%} \right]$ is a matrix of size $[N_{ROI}\times(N_{cond}-1)]$ containing the electric fields of the $N_{cond}-1$ remaining E-MEP curves at 1% stimulator intensity; $\left[ I_{c}^{min} \right]$ is a matrix of size $[N_{ROI}\times(N_{cond}-1)]$, where each column consists of the respective minimal stimulator intensity $I_{c}^{min}$ (the same for each row); $\mathbf{e}_{ref}^{1\%}$ is a vector of size $[N_{ROI}\times1$] containing the electric field of the condition corresponding to the reference I/O curve at 1% stimulator intensity; $\mathbf{I}_{c}^{min}$ and $\mathbf{I}_{c}^{max}$ are two vectors of size $[(N_{conds}-1)\times1]$ containing the minimal and maximal values of the remaining I/O curves; the operator $\circ$ is the Hadamard product, i.e. the element-wise multiplication of the matrix entries; the operator $\otimes$ is the outer product between two vectors resulting in a matrix; and $⊘$ denotes the Hadamard division.

After determining all initial shifts $\tau_{init}$, the optimal shifts $\tau_{opt}$ are determined in the shift space, which has to be defined first. The step size $\Delta E$ is determined by the number of samples $n_{s}$ of the reference curve:

$\Delta E=\frac{e_{ref}^{max}-e_{ref}^{min}}{n_{s}}$	(2)

The electric field simulations are performed assuming a stimulator intensity of 1%. The corresponding electric field is denoted by $e_{ref}^{1\%}$. Due to the linear dependency between the electric field and the stimulator intensity, one can reformulate eq. (2) in terms of stimulator intensity:

$\Delta E=e_{ref}^{1\%}\frac{I_{ref}^{max}-I_{ref}^{min}}{n_{s}}$	(3)

In order to define the shift space, it is necessary to determine the number of steps before and after the step-wise shift procedure $n_{s}^{'}$ and $n_{s}^{''}$, respectively:

$n_{s}^{'}=\frac{e_{ref}^{min}-e_{c}^{min'}}{\Delta E}$	(4)

$n_{s}^{''}=\frac{e_{c}^{max''}-e_{ref}^{max}}{\Delta E}$	(5)

The start point of the transformed I/O curve $e_{c}^{min'}$ (left dotted I/O curve in Fig. S1) is given by:

$e_{ref}^{min}=e_{ref}^{1\%}I_{ref}^{min}=e_{c}^{'1\%}I_{c}^{max} \to e_{c}^{'1\%}=e_{ref}^{1\%}\frac{I_{ref}^{min}}{I_{c}^{max}}$	(6)

$e_{c}^{min'}=e_{c}^{'1\%}I_{c}^{min}$	(7)

Where $e_{c}^{'1\%}$ denotes the fictitious electric field value, the transformed I/O curve would correspond to the situation in which the curve is located at the initial position of the shift space. Inserting eq. (6) into (7) yields:

$e_{c}^{min'}=e_{ref}^{1\%}I_{ref}^{min}\frac{I_{c}^{min}}{I_{c}^{max}}$	(8)

The end point of the transformed I/O curve after it is shifted $e_{c}^{max''}$ (right dotted I/O curve in Fig. S1) is given by:

$e_{ref}^{max}=e_{ref}^{1\%}I_{ref}^{max}=e_{c}^{''1\%}I_{c}^{min} \to e_{c}^{''1\%}=e_{ref}^{1\%}\frac{I_{ref}^{max}}{I_{c}^{min}}$	(9)

$e_{c}^{max''}=e_{c}^{''1\%}I_{c}^{max}$	(10)

Where $e_{c}^{''1\%}$ denotes the fictitious electric field value, the I/O curve would correspond to the situation in which the curve is located at the end point after a complete shift. Inserting eq. (9) into (10) yields:

$e_{c}^{max''}=e_{ref}^{1\%}I_{ref}^{max}\frac{I_{c}^{max}}{I_{c}^{min}}$	(11)

The number of steps before and after the step-wise shift procedure $n_{s}^{'}$ and $n_{s}^{''}$ can be determined by inserting eq. (7) into (4) and eq. (11) into (5).

$n_{s}^{'}=\frac{1 - \frac{I_{c}^{min}}{I_{c}^{max}}}{\frac{I_{ref}^{max}}{I_{ref}^{min}} - 1}n_{s}$	(12)

$n_{s}^{''}=\frac{\frac{I_{c}^{max}}{I_{c}^{min}} - 1}{1 - \frac{I_{ref}^{min}}{I_{ref}^{max}}}n_{s}$	(13)

Next, the shift spaces are constructed and the I/O curves are evaluated on the defined grids. The curves are successively shifted and the optimal displacement $\tau_{opt}$ is determined, where the I/O curves are overlapping optimally. This is done for every of the $N_{conds}-1$ I/O curves excluding the reference. The optimal shifts are stored in a vector $\boldsymbol{\tau}_{opt}$ of size $[(N_{conds}-1)\times1]$.

Finally, the total shifts stored in the matrix $[\tau]$, are determined by:

$[\tau]=[\tau_{init}]+[\tau_{opt}]$,	(14)

where $[\tau_{opt}]$ is a matrix of size $[N_{ROI}\times(N_{cond}-1)]$ containing the optimal shifts $\boldsymbol{\tau}_{opt}$ in each row.

**1.3 Parametrizing the fractional anisotropy of the electrical conductivity tensors**

Here, we present the methodology to vary the level of anisotropy of the electrical conductivity of the head model, which affects the induced electric field profile. The anisotropic properties of GM and WM result from the alignment of the pyramidal cells in the cortex and the fiber pathways in the white matter^[1]^. The level of anisotropy is quantified using the fractional anisotropy $FA$:

$$FA=\sqrt{\frac{\left( \lambda_{1}-\lambda_{2} \right)^{2}+\left( \lambda_{2}-\lambda_{3} \right)^{2}+\left( \lambda_{3}-\lambda_{1} \right)^{2}}{2\left( \lambda_{1}^{2}+\lambda_{2}^{2}+\lambda_{3}^{2} \right)}}$$

(15)

Its three pairwise perpendicular axes of symmetry are given by the eigenvalues $\lambda_{i}$ and eigenvectors $\boldsymbol{v}_{i}$ determined by an Eigenvalue decomposition of the conductivity tensor $\left[ \boldsymbol{\sigma} \right]$:

$$\left[ \boldsymbol{\sigma} \right]=\left[ \mathbf{V} \right]\left[ \boldsymbol{\Lambda} \right]\left[ \mathbf{V}^{-1} \right].$$

(16)

The columns of the matrix $\left[ \mathbf{V} \right]=\left[ \mathbf{v}_{1},\mathbf{v}_{2},\mathbf{v}_{3} \right]$ contain the eigenvectors $\mathbf{v}_{i}$ and the diagonal matrix $\left[ \boldsymbol{\Lambda} \right]$ the corresponding eigenvalues $\lambda_{i}$. The $FA$ of isotropic tissues is 0, or 1 in case of completely restricted tensors. Every tetrahedron in GM and WM obeys its own conductivity tensor and hence $FA$ parameter. Varying the $FA$ for every single element independently is not practical and would vastly increase the number of random variables making the problem intractable. Hence, in order to incorporate the level of anisotropy into the uncertainty analysis of the TMS problem, an anisotropy variation parameter $\alpha$ is introduced. This parameter is aimed to control the level of anisotropy in the whole brain model at once. A tensor can be visualized as an ellipsoid as shown in Fig. S1(a). The anisotropy parameter is defined similar as the $FA$, i.e. $\alpha=\left[ 0, 1 \right]=\left\{ \alpha\mathbb{\in R|}0\leq\alpha\leq1 \right\}$. It is initialized at a value of $\alpha=0.5$ for every tensor. Decreasing $\alpha$ towards 0 morphs the ellipsoid into a sphere, whereas increasing it towards 1 makes the tensor more restrictive in the direction of its major axis. In order to perform the transformation of the tensor, the ellipsoid is subdivided into three ellipses lying orthogonal to each other in its three principal planes. Each ellipse has an eccentricity $e_{i,j}$ given by:

|  | $e_{1,2}=\sqrt{1-\left( \frac{\lambda_{2}}{\lambda_{1}} \right)^{2}}$ | $e_{1,3}=\sqrt{1-\left( \frac{\lambda_{3}}{\lambda_{1}} \right)^{2}}$ | $e_{2,3}=\sqrt{1-\left( \frac{\lambda_{3}}{\lambda_{2}} \right)^{2}}$ | (17) |
| --- | --- | --- | --- | --- |

The eccentricities $e_{i,j}$ are transformed to the normalized $\alpha$ scale. By changing $\alpha$, the three eccentricities $e_{i,j}$ are modified in a linear sense as it is depicted in Fig. S1(b). The transformed eccentricities $\tilde{e}_{i,j}$ are given by:

$$\tilde{e}_{i,j}=\left\{ \begin{matrix} 2\alpha e_{i,j}, \\ 2\alpha\left( 1-e_{i,j} \right)+2e_{i,j}-1, \end{matrix} \begin{matrix} \mathrm{if} \alpha\leq0.5 \\ \mathrm{if} 0.5<\alpha\leq1 \end{matrix} \right.$$

(18)

The scaled eigenvalues $\tilde{\lambda}_{i}$ are normalized such that the volume of the unscaled and scaled conductivity tensors are equal:

|  | $\tilde{\lambda}_{1}=k_{1}\left( \prod_{i=1}^{3} \frac{\lambda_{i}}{k_{i}} \right)^{\frac{1}{3}}$ | $\tilde{\lambda}_{2}=k_{2}\left( \prod_{i=1}^{3} \frac{\lambda_{i}}{k_{i}} \right)^{\frac{1}{3}}$ | $\tilde{\lambda}_{3}=k_{3}\left( \prod_{i=1}^{3} \frac{\lambda_{i}}{k_{i}} \right)^{\frac{1}{3}}$ | (19) |
| --- | --- | --- | --- | --- |

with

|  | $k_{1}=1$ | $k_{2}=\sqrt{1-\tilde{e}_{1,2}^{2}}$ | $k_{3}=\sqrt{1-\tilde{e}_{1,3}^{2}}$ | (20) |
| --- | --- | --- | --- | --- |

Since the principal axes of the conductivity tensor in $\left[ \mathbf{V} \right]$ are not affected by the transformation, the transformed diffusion tensor $\left[ \tilde{\boldsymbol{\sigma}} \right]$ can be determined by:

$$\left[ \tilde{\boldsymbol{\sigma}} \right]=\left[ \mathbf{V} \right]\left[ \tilde{\boldsymbol{\Lambda}} \right]\left[ \mathbf{V}^{-1} \right]$$

(21)

where $\left[ \tilde{\boldsymbol{\Lambda}} \right]$ contains the scaled eigenvalues from (16) in its diagonal. The anisotropy scaling parameter $\alpha$ is included in the uncertainty analysis and modelled as a beta distributed random variable with the parametrization given in Table 1.

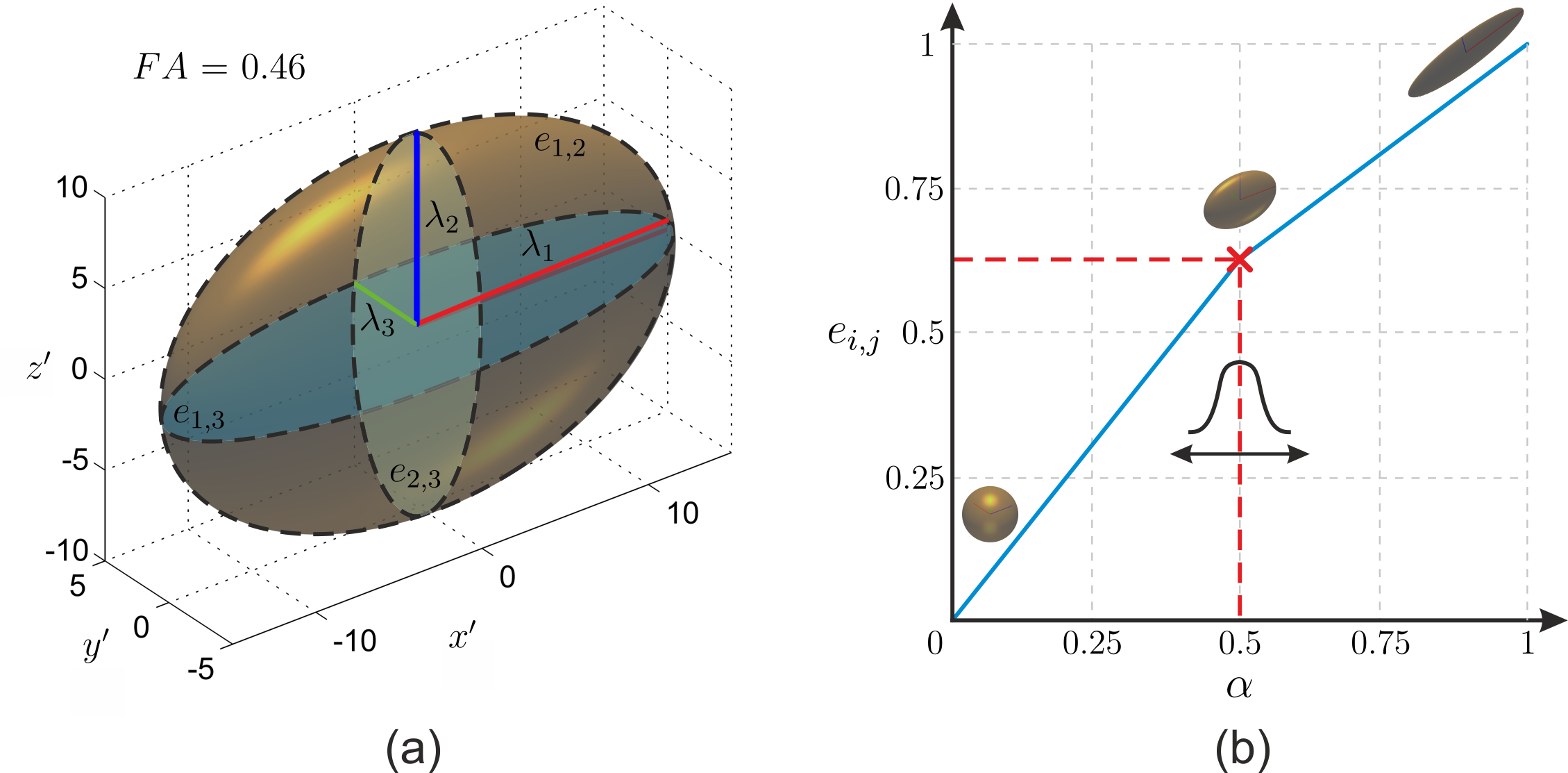

**Figure S2:** (a) Exemplary conductivity tensor with a fractional anisotropy of $FA=0.46$ determined from diffusion weighted magnetic resonance images considering a volume normalized approach (Güllmar et al., 2010). Highlighted are the eigenvectors, the corresponding eigenvalues $\lambda_{i}$¸ and the three ellipses obeying the eccentricities $e_{i,j}$ lying in the $i,j$-planes; (b) Anisotropy parameter $\alpha$ scaling the fractional anisotropy $FA$ and eccentricities $e_{i,j}$ of the conductivity tensor. The original shape of the tensor is preserved for $\alpha=0.5$.

**1.4 Nonintrusive Generalized Polynomial Chaos**

When using nonintrusive generalized polynomial chaos (gPC), the whole congruence factor calculation scheme is treated as a black box system. In the present context, its inputs are the random variables $\boldsymbol{p}$, i.e., the uncertain parameters of the model, and the output is the position wise congruence factor $c\left( \boldsymbol{r},\boldsymbol{p} \right)$. The gPC is based on the polynomial approximation of the output quantity:

$$c\left( \boldsymbol{r},\boldsymbol{p} \right)=\sum_{\boldsymbol{\alpha}\in\text{A}} u_{\boldsymbol{\alpha}}\left( \boldsymbol{r} \right)\Psi_{\boldsymbol{\alpha}}\left( \boldsymbol{p} \right)$$

(22)

where $\Psi_{\alpha}\left( \boldsymbol{p} \right)=\prod_{i=1}^{d} \psi_{\alpha_{i}}\left( \boldsymbol{p} \right)$ are the joint polynomial basis functions of the gPC, which are composed of separately defined polynomials $\psi_{\alpha_{i}}\left( \boldsymbol{p} \right)$ for each random variable. The polynomial families are chosen to be orthogonal in the normed space induced by the probability density functions (Askey and Wilson, 1985). The multi-index $\boldsymbol{\alpha}$ includes the degrees of the individual polynomials. The central part of the algorithm is the calculation of the coefficients $u_{\boldsymbol{\alpha}}\left( \boldsymbol{r} \right)$ of the polynomials at each cortical location $\boldsymbol{r}$. Reformulating (22) in matrix form, the gPC coefficients are estimated using the method of least squares:

$$\left[ \mathbf{U} \right]=\left[ \boldsymbol{\Psi} \right]^{+}\left[ \mathbf{C} \right]$$

(23)

where $\left[ \mathbf{U} \right]$ is the coefficient matrix of size $\left[ N_{u}\times N_{ROI} \right]$, whose columns are the $N_{u}$ gPC-coefficients for each of the $N_{ROI}$ congruence factors. $\left[ \boldsymbol{\Psi} \right]$ is the gPC-matrix of size $\left[ N_{g}\times N_{u} \right]$, whose columns are the polynomial basis functions evaluated at the grid points on a grid *G* over the parameter space (requiring $N_{g}$ congruence factor calculations); the superscript + indicates the pseudo-inverse; and $\left[ \mathbf{C} \right]$ is the congruence factor matrix of size $\left[ N_{g}\times N_{ROI} \right]$, whose columns are the model solutions of the $N_{ROI}$ congruence factors, i.e. finite elements, in the ROI. An adaptive algorithm is used, which consecutively extends the gPC matrix $\left[ \boldsymbol{\Psi} \right]$ by adding polynomials to the active set while also enlarging the grid *G* to solve the least square problem (Saturnino et al., 2018^[2]^). The ratio between the number of forward solutions and the number of polynomials is set to ${N_{g}}/{N_{ROI}}=1.5$, thus ensuring a sufficient degree of overdeterminedness.

After determining the gPC-coefficient matrix $\left[ \mathbf{C} \right],$ determination of the stochastic properties and sensitivity analysis of the congruence factors can be conducted with high computational efficacy in a post-processing step. The expectation $\mu\left( \boldsymbol{r} \right)$ and variance $\nu\left( \boldsymbol{r} \right)$ of the congruence factors $c\left( \boldsymbol{r} \right)$ can be directly calculated from the gPC-coefficients:

$$\mu\left( \boldsymbol{r} \right)=u_{\alpha_{0}}\left( \boldsymbol{r} \right)$$

(24)

$$\nu\left( \boldsymbol{r} \right)=\sum_{\boldsymbol{\alpha\in}\text{A \textbackslash}\boldsymbol{\alpha}_{0}} u_{\alpha}^{2}\left( \boldsymbol{r} \right)$$

(25)

The elementwise relative standard deviation (*RSD*) is then given by the ratio of the standard deviation and the expectation, i.e. $RSD={\sqrt{\nu\left( \boldsymbol{r} \right)}}/{\mu\left( \boldsymbol{r} \right)}$. For the sensitivity analysis, the Sobol indices $S_{i}\left( \boldsymbol{r} \right)$ are calculated. They represent portions of the total variance $\nu\left( \boldsymbol{r} \right)$, which are due to individual parameters $p_{i}$ or a combination thereof [Sobol (2001), Sudret (2008)]. For each $S_{i}\left( \boldsymbol{r} \right)$, a separate subset of multi-indices $\text{A}_{i}\subseteq\text{A}$ is constructed that references those polynomials that depend on the particular parameter combination of interest, e.g. the conductivity of GM or the combination between different measurement parameters. The Sobol index $S_{i}\left( \boldsymbol{r} \right)$ is calculated similarly to the variance (25), but over the reduced set $\text{A}_{i}$:

$$S_{i}\left( \boldsymbol{r} \right)=\frac{1}{\nu\left( \boldsymbol{r} \right)}\sum_{\alpha\in\text{A}_{i}} u_{\alpha}^{2}\left( \boldsymbol{r} \right)$$

(26)

**1.4 Model to conduct the uncertainty and sensitivity analysis of the congruence factor**

The model to determine the congruence factor is shown in Fig. S2. The input parameters of the model are the electrical conductivities of GM, WM, and CSF $[\boldsymbol{\sigma}]$, the anisotropy scaling parameter $\alpha$, and the fitted MEP curve parameters $x_{0,i}$**.** By varying the electrical conductivities and the level of anisotropy, the electric field of each experimental condition would have to be recomputed in each calculation step of the congruence factor, requiring $N_{c}$ costly FEM simulations in each iteration. In order to decrease the computational cost substantially, the electric fields are estimated based on a separate gPC-approximation. The electric field gPCs are determined with the adaptive algorithm proposed by Saturnino et al., (2018)^3^ in a pre-processing step with a leave-one-out cross validation error of <0.05%. The electric field gPC approximations act as sub-modules in the model of the congruence factor. Depending on the experimental condition, the electric field gPCs converged after 138 simulations using a maximum polynomial approximation order of 4 considering a maximum interaction order of 3. The E-MEP curves are constructed by combining the electric fields with the measured I/O-curves. This is the input data to determine the congruence factor in each element in the ROI.

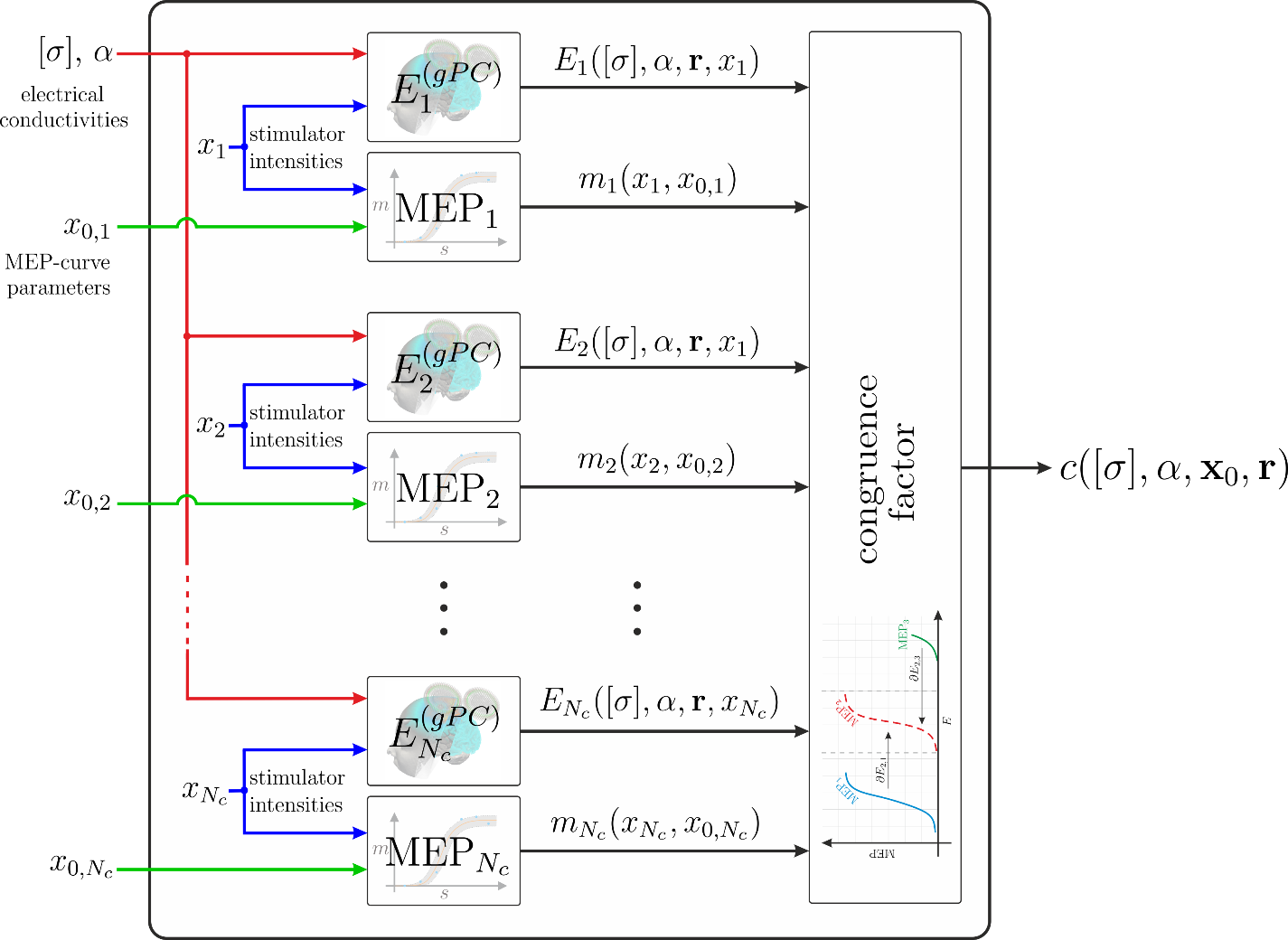

**Figure S3:** Model of the congruence factor used in the gPC based uncertainty and sensitivity analysis. The electrical conductivities of GM, WM, and CSF $[\boldsymbol{\sigma}]$, the anisotropy scaling parameter $\alpha$, and the fitted MEP curve parameters $x_{0,i}$ are modelled as beta distributed random variables. The electric fields of the individual conditions $E_{i}^{(gPC)}$ are computed based on a separate gPC approximation to avoid $N_{c}$ FEM calculations in every calculation step of the congruence factor and hence to decrease the computational cost substantially. The electric field gPCs are determined in a pre-processing step considering a relative error bound of $<0.05\%$.

**1.5 Optimization of TMS coil position and orientation**

We defined the individual congruence factor hotspot as a cortical target and projected it to the skin surface. Subsequently, we calculated a dense grid of coil positions and orientations as shown in Fig. S3. We were searching for the optimal coil position and orientation in a radius of 20 mm around the projected point. The distance between the tested coil positions is about 1.5 mm. Their orientation was defined in the interval from -60° to 60° around the PA-45 orientation in steps of 15°. This resulted in about 4500 different coil positions and orientations for which we determined the magnitude of the electric field in the predefined target using SimNIBS. Subsequently, we selected the optimal coil position and orientation maximizing the electric field magnitude at the cortical target. We implemented the optimization routine in SimNIBS and it will be published in an upcoming release.

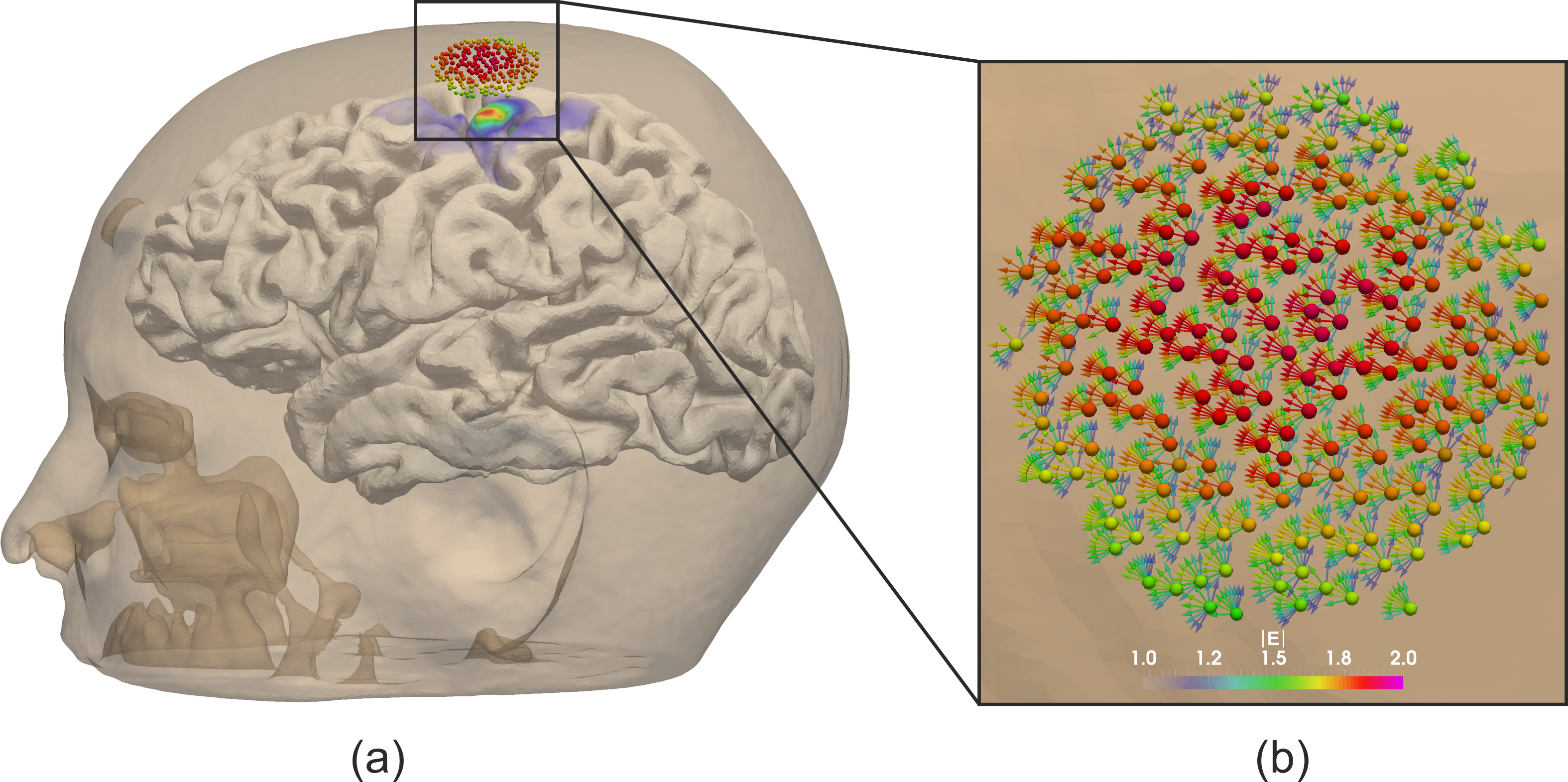

**Figure S4:** Coil positions and orientations used in the optimization study. For every coil position and orientation, the electric field magnitude was determined in the congruence factor hotspot (cortical target). The color indicates the electric field magnitude at the cortical target.

**2. RESULTS**

**2.1 Coil positions and orientations of Experiment II, subject 15**

For the validation study, we repeated Experiment II for subject 15 from group III (c.f. Fig. 7). For this subject however, we had to increase the number of conditions to 30, due to the lack of a single, pronounced congruence factor hotspot using the first 20 conditions. We observed high similarity between the I/O curves and the induced electric field profiles at various cortical positions in this subject. To increase the electric field variance, we added further conditions at different positions, orientations and tilting angles of the TMS coil. The resulting coil positions and orientations are shown in Fig. S4.

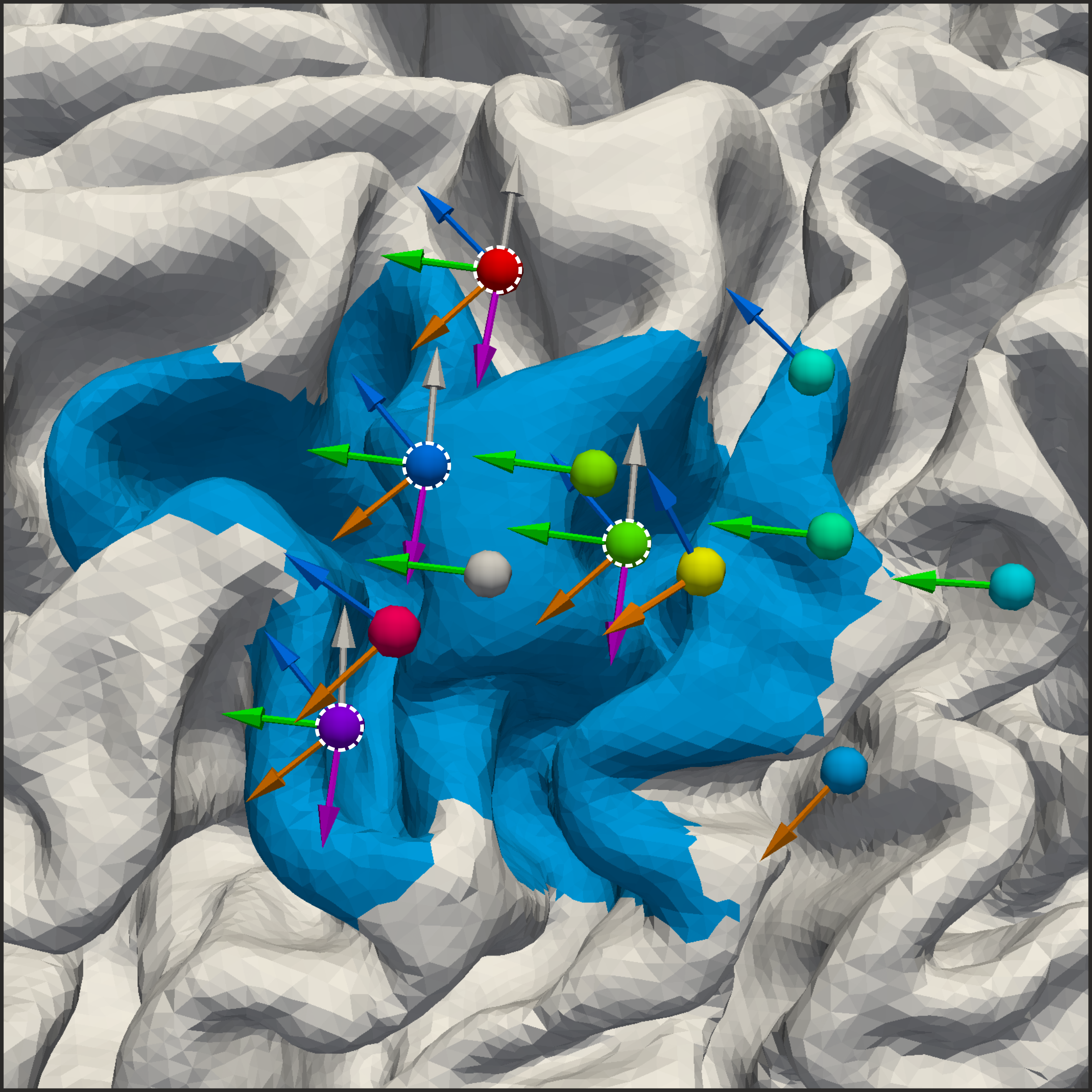

**Figure S5:** Coil positions and orientations used to determine the site of stimulation for subject 15 in Experiment II. The 20 coil positions, which were sufficient for the other subjects are highlighted with a dotted white circle. The additional coil positions and orientations lead to an increase in electric field variability in the ROI, which was necessary to determine the congruence factor hotspot successfully.

**2.2 Permutation study**

The results of the permutation study for subjects S8 and S12 are shown in Fig. S3 and Fig. S4, respectively. The hotspot areas in Fig. S3(b) and Fig. S4(b) show very similar convergence across subjects regarding the number of conditions $k$ and so do the cross-correlations of the electric fields (Fig. S3(c) and S4(c)). For all subjects, it can be observed that condition combinations of electric fields with low cross-correlation are preferable to determine focal congruence factor maps (black lines in Fig. S3(b), S4(b), S3(c), and S4(c)). The chord graphs in Fig. S3(d) and Fig. S4(d) show that stimulation sites around M1 together with different coil orientations are preferable to determine the site of stimulation.

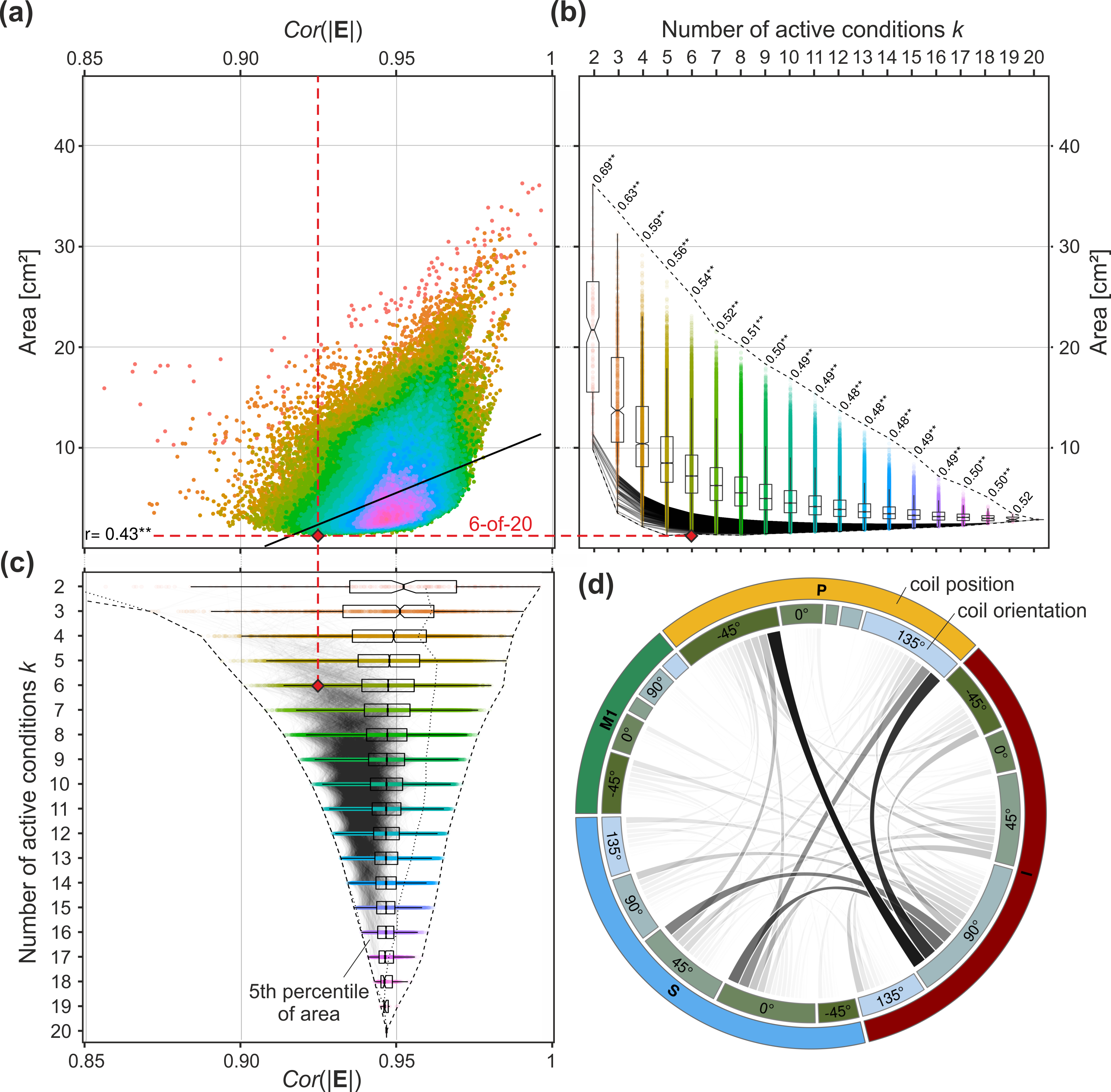

**Figure S6:** Permutation analysis. Interrelating the cross-correlation of the electrical fields from the different experimental conditions of Experiment II for subject S8 with the corresponding hotspot area. The hotspot area is defined as the region where $c>30$. For each case $k$, the congruence factor is determined $\left( \begin{matrix} 20 \\ k \end{matrix} \right)$times. (a) Relationship between the cross-correlation of the electric field magnitude and the resulting hotspot area size. Colors: active conditions $k$. The correlation coefficient between the hotspot size and the cross-correlation of the electric fields over all $k$ (black line) is $r=0.43 (p\ll.001)$. (b) Boxplot of the hotspot area of the congruence factor in dependence of the number of active conditions $k$. Box areas indicate the 25% to 75% quantiles with notch at median. Correlation coefficients between the hotspot size and the cross-correlation of the electric fields for each *k* are given. ** depict $p<.01$ (after Bonferroni correction). Grey lines: 5^th^ percentile of the best condition combinations for each *k* (same as in (c)). Dashed lines: absolute range. The variation of the hotspot area size decreases with increasing $k$. (c) Relationship between the cross-correlation coefficient of the electric fields and the number of active conditions $k$. Box areas indicate the 25% to 75% quantiles with notch at median. Grey lines: 5^th^ percentile of the best condition combinations for each $k$ (same as in (b)). Dashed lines: absolute range. The dashed red lines highlight the case of the 6 coil positions and orientations, where the congruence factor map was most focal. Its cross-correlation coefficient is with 0.925 lower than the first quartile of possible solutions. (d) Chord graph highlighting the interaction and relative contribution between different coil positions (outer circle) and coil orientations (inner circle) from the 5^th^ percentile of best condition combinations over all $k$ resulting in small hotspot areas (highlighted with black lines in (b)).

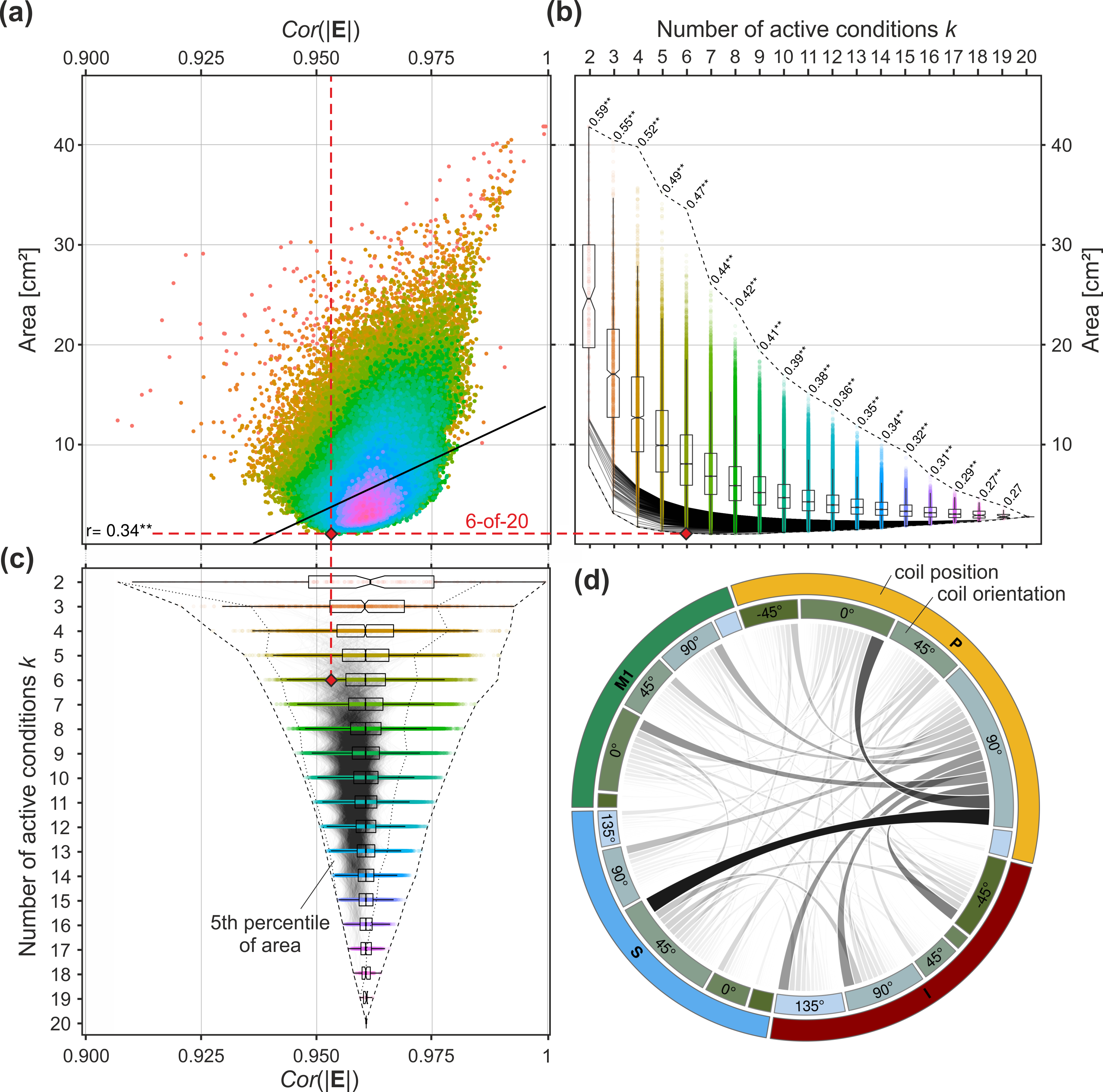

**Figure S7:** Permutation analysis. Interrelating the cross-correlation of the electrical fields from the different experimental conditions of Experiment II for subject S12 with the corresponding hotspot area. The hotspot area is defined as the region where $c>30$. For each case $k$, the congruence factor is determined $\left( \begin{matrix} 20 \\ k \end{matrix} \right)$times. (a) Relationship between the cross-correlation of the electric field magnitude and the resulting hotspot area size. Colors: active conditions $k$. The correlation coefficient between the hotspot size and the cross-correlation of the electric fields over all $k$ (black line) is $r=0.34 (p\ll.001)$. (b) Boxplot of the hotspot area of the congruence factor in dependence of the number of active conditions $k$. Box areas indicate the 25% to 75% quantiles with notch at median. Correlation coefficients between the hotspot size and the cross-correlation of the electric fields for each *k* are given. ** depict $p<.01$ (after Bonferroni correction). Grey lines: 5^th^ percentile of the best condition combinations for each *k* (same as in (c)). Dashed lines: absolute range. The variation of the hotspot area size decreases with increasing $k$. (c) Relationship between the cross-correlation coefficient of the electric fields and the number of active conditions $k$. Box areas indicate the 25% to 75% quantiles with notch at median. Grey lines: 5^th^ percentile of the best condition combinations for each $k$ (same as in (b)). Dashed lines: absolute range. The dashed red lines highlight the case of the 6 coil positions and orientations, where the congruence factor map was most focal. Its cross-correlation coefficient is with 0.953 lower than the first quartile of possible solutions. (d) Chord graph highlighting the interaction and relative contribution between different coil positions (outer circle) and coil orientations (inner circle) from the 5^th^ percentile of best condition combinations over all $k$ resulting in small hotspot areas (highlighted with black lines in (b)).

**2.2 Uncertainty and sensitivity analysis**

The uncertainty analysis of the congruence factor was also conducted for subjects S8 and S12 considering the best 6-of-20 results of Experiment II. The uncertainties of the MEP turning point parameters are shown in Table S1 and S2 for subject S1 and S12, respectively.

The results of the uncertainty and sensitivity analysis of subject S8 and S12 are shown in Fig. S5 and Fig. S6, respectively. The relative average sensitivity of the congruence factor with respect to the measured I/O curves is 40%, 12%, and 49.9% for subjects S1, S8, and S12, respectively. The difference of its relative contribution in subject S8 originates from the fact that the I/O curve parameters have an overall lower uncertainty.

**Table S1:** *Limits of the MEP turning point parameters for subject S8,*

| Parameter | Min | Max |
| --- | --- | --- |
| $x_{0,S\left( 0^{\circ} \right)}$ | 74.6 [A/µs] | 78.5 [A/µs] |
| $x_{0,S\left( 45^{\circ} \right)}$ | 74.5 [A/µs] | 77.1 [A/µs] |
| $x_{0,S\left( 135^{\circ} \right)}$ | 84.9 [A/µs] | 88.7 [A/µs] |
| $x_{0,I\left( -45^{\circ} \right)}$ | 108.8 [A/µs] | 120.3 [A/µs] |
| $x_{0,I\left( 90^{\circ} \right)}$ | 92.8 [A/µs] | 112.8 [A/µs] |
| $x_{0,P\left( 135^{\circ} \right)}$ | 111.1 [A/µs] | 122.8 [A/µs] |

*Notes: These values were considered in the uncertainty and sensitivity analysis to determine the site of stimulation by means of the congruence factor. All parameters are modeled as symmetric bell shaped* $\beta$*-distributions with shape* parameters $p=q=4$.

**Table S2:** *Limits of the MEP turning point parameters for subject S12*

| Parameter | Min | Max |
| --- | --- | --- |
| $x_{0,S\left( 90^{\circ} \right)}$ | 70.3 [A/µs] | 88.3 [A/µs] |
| $x_{0,I\left( -45^{\circ} \right)}$ | 115.8 [A/µs] | 131.8 [A/µs] |
| $x_{0,I\left( 90^{\circ} \right)}$ | 95.8 [A/µs] | 101.8 [A/µs] |
| $x_{0,P\left( 0^{\circ} \right)}$ | 83.7 [A/µs] | 92.5 [A/µs] |
| $x_{0,P\left( 45^{\circ} \right)}$ | 90.9 [A/µs] | 120.9 [A/µs] |
| $x_{0,P\left( 90^{\circ} \right)}$ | 110.5 [A/µs] | 136.5 [A/µs] |

*Notes: which are considered in the uncertainty and sensitivity analysis to determine the site of stimulation by means of the congruence factor. All parameters are modeled as symmetric bell shaped* $\beta$*-distributions with shape parameters* $p=q=4$*.*

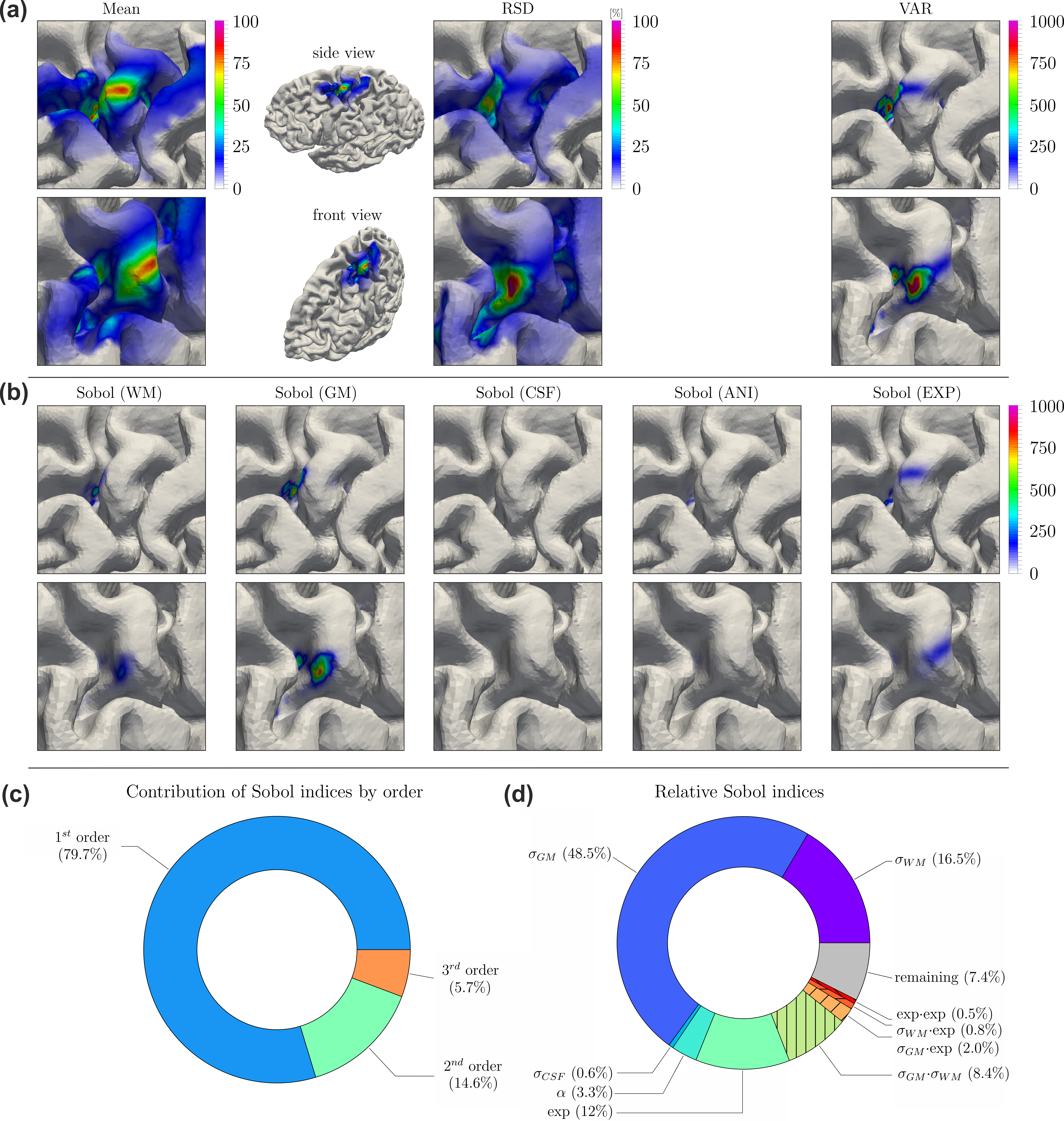

**Figure S8:** Uncertainty and sensitivity analysis of the congruence factor determined for subject S8 in the 6-of-20 analysis of Experiment II. (a) Expected value (Mean), relative standard deviation (RSD), and variance (VAR) of the congruence factor. (b) Absolute first order Sobol indices. The normalization with respect to the total variance in (12) was avoided to strengthen the focus on regions of high variance. The Sobol index maps of the individual MEP parameters resulting from uncertainties in the experimental data are summarized into one Sobol index map “Sobol (EXP)”. (c) Relative contribution of the average Sobol indices by order. (d) Average composition of the first order Sobol indices. For (a) and (b) two different perspectives are shown (top and bottoms rows), in order to improve visibility of the effects.

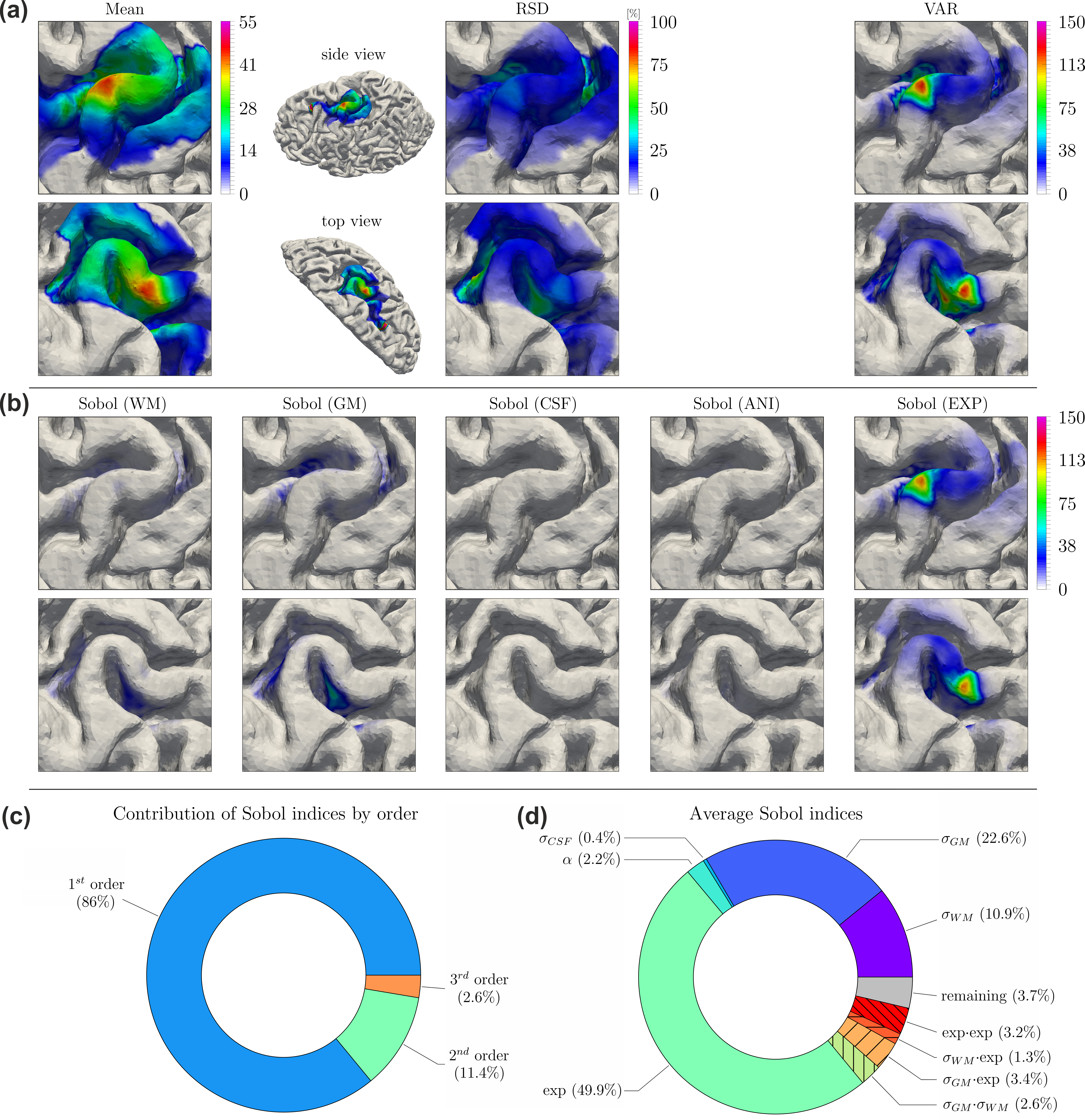

**Figure S9:** Uncertainty and sensitivity analysis of the congruence factor determined for subject S12 in the 6-of-20 analysis of Experiment II. (a) Expected value (Mean), relative standard deviation (RSD), and variance (VAR) of the congruence factor. (b) Absolute first order Sobol indices. The normalization with respect to the total variance in (12) was avoided to strengthen the focus on regions of high variance. The Sobol index maps of the individual MEP parameters resulting from uncertainties in the experimental data are summarized into one Sobol index map “Sobol (EXP)”. (c) Relative contribution of the average Sobol indices by order. (d) Average composition of the first order Sobol indices. For (a) and (b) two different perspectives are shown (top and bottoms rows), in order to improve visibility of the effects.

1. Kandel, ER, Schwartz JH, Jessell T, Siegelbaum SA, Hudspeth AJ, Mack S, Principles of neural science. Fifth edition, 2013, New York, Lisbon, London: McGraw-Hill Medical.
2. Saturnino GB, Thielscher A, Madsen KH, Knösche TR, Weise K, A principled approach to conductivity uncertainty analysis in electric field calculations. *NeuroImage*, 2018, in press. (<https://doi.org/10.1016/j.neuroimage.2018.12.053>)
